# TBK1 phosphorylation activates LIR-dependent degradation of the inflammation repressor TNIP1

**DOI:** 10.1101/2022.03.02.482646

**Authors:** Jianwen Zhou, Nikoline Lander Rasmussen, Hallvard Lauritz Olsvik, Vyacheslav Akimov, Zehan Hu, Gry Evjen, Blagoy Blagoev, Trond Lamark, Terje Johansen, Jörn Dengjel

## Abstract

Limitation of excessive inflammation due to selective degradation of pro-inflammatory proteins is one of the cytoprotective functions attributed to autophagy. In the current study, we highlight that selective autophagy also plays a vital role in promoting the establishment of a robust inflammatory response. Under inflammatory conditions, here TLR3-activation by poly(I:C) treatment, the inflammation repressor TNIP1 (TNFAIP3 interacting protein 1) is phosphorylated by TBK1 (Tank-binding kinase 1) activating a LIR motif that leads to the selective autophagy-dependent degradation of TNIP1, supporting expression of pro-inflammatory genes and proteins. Thus, similarly as in cancer, autophagy may play a dual role in controlling inflammation depending on the exact state and timing of the inflammatory response.

**Summary:** Autophagy is well known for its anti-inflammatory effects. Here, we highlight that selective, autophagy-dependent degradation of the inflammation repressor TNIP1 supports pro-inflammatory gene and protein expression. Similarly as in cancer, autophagy appears to play a dual role in controlling inflammation.

## Introduction

Macroautophagy, hereafter referred to as autophagy, is a primarily cytoprotective process that leads to the removal and lysosomal degradation of non-functional and/or superfluous cytoplasmic material. Autophagy is a constitutive process but may also be triggered by various forms of cell stress (Mizushima and Levine, 2020). It involves the formation of a double membrane vesicle known as autophagosome that enwraps portions of the cytoplasm, including organelles. The content of the autophagosome is then targeted for degradation via the lysosome. Constitutive autophagy is regarded as a non-selective, bulk process, whereas stress-induced autophagy is often selective, aiming at removing the causes of stress, e.g. depolarized mitochondria in oxidative stress (Youle, 2019). Selective autophagy is carried out either by direct interactions between cargo and lipidated human ATG8 family members (MAP1LC3-A, −B, −C, GABARAP, GABARAPL1, GABARAPL2, commonly referred to as LC3s) that are anchored to autophagosomal membranes, or by indirect interactions in which so-called selective autophagy receptors (SARs) tether cargo to LC3s (Johansen and Lamark, 2020). Cargo and receptors are then both degraded within lysosomes (Morishita and Mizushima, 2019). p62/SQSTM1, which is considered the founding member of the protein class of p62/SQSTM1- like receptors (SLRs), recognizes poly-ubiquitinated proteins/organelles destined for lysosomal degradation via its ubiquitin-associated (UBA) domain and interacts with lipidated LC3s via its LC3-interacting region (LIR) motif (Johansen and Lamark, 2020; Pankiv et al., 2007). SLRs are involved in the selective, autophagy-dependent degradation of a highly diverse set of substrates (Zellner et al., 2021).

The limitation of deleterious inflammatory responses is one of the cytoprotective functions attributed to autophagy (Deretic and Levine, 2018). The link between autophagy and tissue inflammation became obvious by genome-wide association studies that identified *ATG16L1* as susceptibility locus for Crohn disease (Hampe et al., 2007). Since then several autophagy loci have been linked to inflammatory and autoimmune disorders (Mizushima and Levine, 2020). Selective degradation of inflammasome components, e.g. by p62/SQSTM1, is among the best understood functions of autophagy in limiting tissue inflammation (Deretic and Levine, 2018; Samie et al., 2018). However, autophagy was also shown to support unconventional secretion of the pro-inflammatory cytokine IL1B (Dupont et al., 2011; Zhang et al., 2015), which argues for a fine-tuned and balanced role of autophagy in regulating inflammatory responses.

TNIP1 (also known as ABIN-1, Naf1 and VAN) is a ubiquitin-binding adaptor protein that has been implicated as a negative regulator of inflammatory signaling and cytokine-induced cell death (Dziedzic et al., 2018; Gao et al., 2011; Oshima et al., 2009; Su et al., 2019; Zhou et al., 2011). Interestingly, a recent study reported the selective autophagy-dependent degradation of TNIP1 via interaction with the SLR optineurin (OPTN), supporting senescence-associated inflammation (Lee et al., 2021). While TNIP1 itself shows no catalytic activity, its ability to bind linear polyubiquitin chains through its Ub-binding domain in ABIN proteins and NEMO (UBAN) is important for its anti-inflammatory function (Nanda et al., 2011; Wagner et al., 2008). A number of studies suggest that TNIP1 excerts its negative function by recruiting the ubiqutitin-editing enzyme TNFAIP3 (also known as A20) to polyubiquitinated targets (Dziedzic et al., 2018; Gao et al., 2011; Mauro et al., 2006). However, the exact mechanisms behind this negative regulation is not completely understood. Furthermore, TNIP1 shows activity independent of TNFAIP3 (Kattah et al., 2018; Oshima et al., 2009). The importance of TNIP1 as an anti-inflammatory signal transducer is highlighted by numerous studies implicating TNIP1 dysregulation in autoimmune disorders (Shamilov and Aneskievich, 2018). Uncovering the molecular function and dynamics of TNIP1 could therefore be valuable in understanding the mechanisms behind such complex disorders.

In the current study, we identify TNIP1 as an autophagy substrate, which is selectively degraded under inflammatory conditions. We highlight that TNIP1 fulfills the structural characteristics of an autophagy receptor with an oligomerization domain, a ubiquitin-binding domain and a LIR motif. Upon TLR3 activation TNIP1 is phosphorylated by TBK1 on LIR proximal serine residues to increase binding to LC3s. Hence, TNIP1 is selectively degraded by autophagy in order to promote a competent initiation of pro-inflammatory signaling.

## Results

### TNIP1 is ubiquitinated and degraded within lysosomes

To identify new proteins potentially involved in autophagy regulation or autophagosomal targeting, we screened ubiquitination dynamics by quantitative mass spectrometry (MS)-based proteomics. U2OS and HeLa cells were differentially labeled by stable isotope labeling by amino acids in cell culture (SILAC) and autophagy was induced by inhibiting MTORC1 by rapamycin. In parallel, lysosomal degradation was blocked by concanamycin A (ConA), an inhibitor of lysosomal V-type ATPase (Klionsky et al., 2021). Respective cell lysates were mixed, proteins digested with the endoprotease LysC and ubiquitinated peptides were enriched using the UbiSite approach followed by MS-based identification and quantification (Akimov et al., 2018b) (Fig 1A). We identified more than 9,000 ubiquitination sites, of which more than 2,000 were quantified in minimally three biological replicates and could be localized clearly to specific amino acid residues (class I sites, Fig 1B, supplemental Table S1) (Olsen et al., 2006). These sites were used for further analyses. Comparing abundance changes of ubiquitination sites of cells treated with rapamycin with cells treated with rapamycin and ConA, 148 sites were identified as significantly regulated, the majority being more abundant in cells in which lysosomal degradation was inhibited by ConA (paired t-test, FDR<0.05, Fig 1C). As anticipated, we identified many proteins involved in autophagosomal biogenesis and target recruitment. Importantly, four central SLRs, p62/SQSTM1, NBR1, CALCOCO2/NDP52, and TAX1BP1, were identified as being increasingly ubiquitinated (Fig 1C), indicating that our experimental strategy was successful (Johansen and Lamark, 2020).

**Figure 1.**
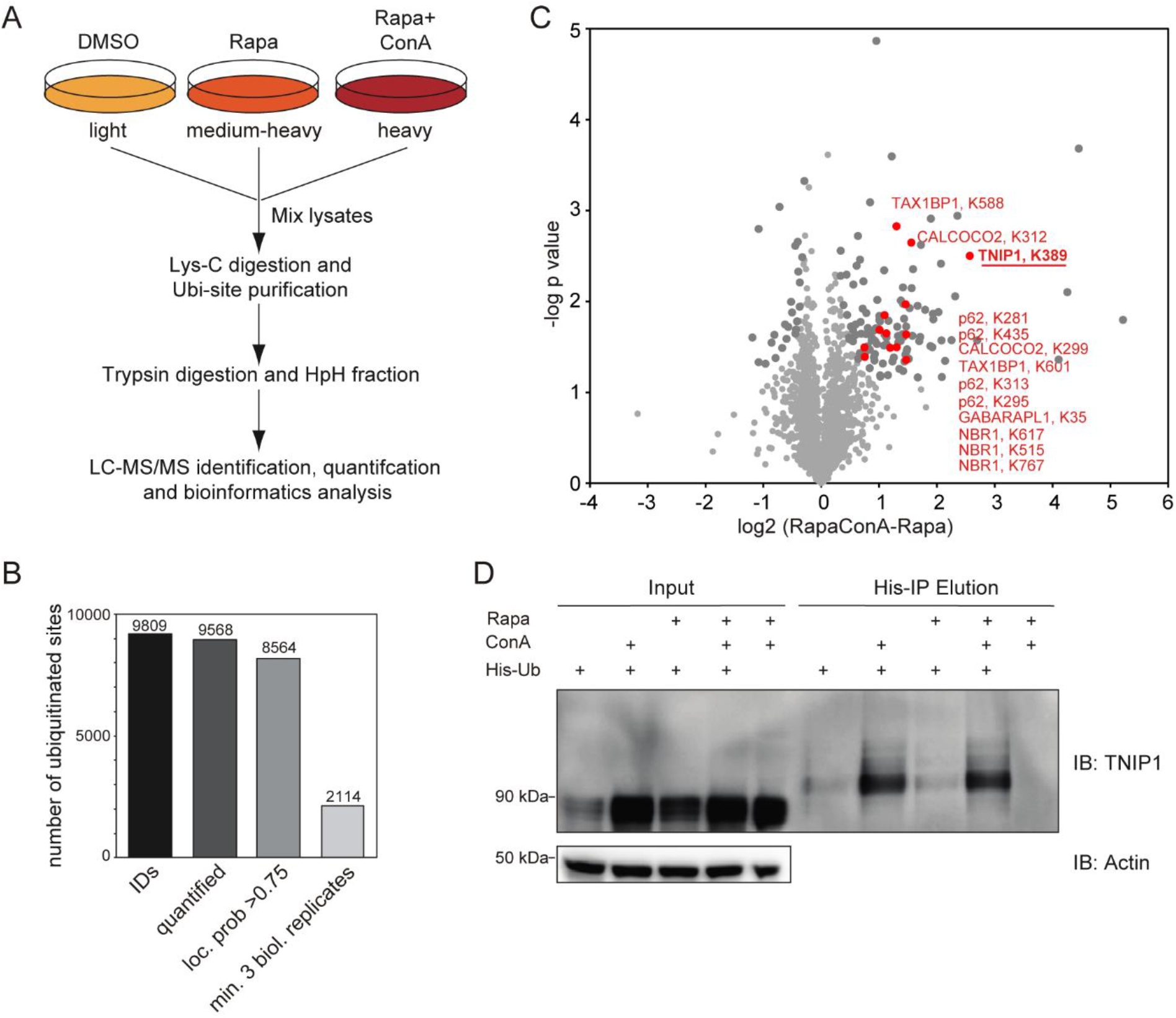
Ubiquitination and lysosomal degradation of TNIP1. A MS workflow to quantify ubiquitination sites potentially involved in autophagy-dependent lysosomal protein degradation. U2OS and HeLa cells were SILAC labeled and treated for 4 h with 100 nM rapamycin (Rapa), Rapa and 2 nM concanamycin A (ConA), or DMSO as control. After mixing of lysates, proteins were digested with Lys-C endoproteinase and ubiquitinated peptides were enriched using the UbiSite approach (Akimov et al., 2018). Enriched peptides were digested with trypsin, followed by high-pH reversed phase fractionation (Batth & Olsen, 2016) and shotgun LC-MS/MS analysis. B Detected ubiquitination sites. In three biological replicates, 9’809 ubiquitination sites were identified of which 9–568 were quantified. C Volcano plot highlighting significantly regulated ubiquitination sites. Significantly regulated sites comparing Rapa with Rapa+ConA treated cells are highlighted in dark grey (paired t-test, FDR<0.05, S0=0.1, 148 sites in total; see supplemental Table S1). Non-regulated sites are colored in light grey. Sites identified on known autophagy receptors are colored in red exemplifying data quality. The newly identified site on TNIP1, K389 is highlighted in bold red. D TNIP1 gets ubiquitinated and degraded in the lysosome. U2-OS-StUbEx cells inducibly expressing His-FLAG-tagged ubiquitin at endogenous levels were used to enrich ubiquitinated proteins (Akimov et al., 2014). Under control conditions as well as under 4 h 100 nM rapamycin treatment TNIP1 got ubiquitinated as shown by anti-TNIP1 immunoblots. Ubiquitinated TNIP1 was stabilized by the addition of concanamycin A indicating its lysosomal degradation in treated and nontreated cells. The same was observed for starved cells (HBSS treatment; see suppl. Fig S1). Actin was used as loading control.

One protein that caught our attention was TNIP1/ABIN-1 that was identified as significantly ubiquitinated on the amino acid residue Lys389 (Fig 1C). TNIP1 is a key repressor of inflammatory signaling (Shamilov and Aneskievich, 2018), and posttranslational mechanisms regulating its protein abundance are largely unknown. In a reverse affinity purification (AP), we used U2OS-StUbEx cells inducibly expressing 6His-FLAG-tagged ubiquitin (Akimov et al., 2014), treated cells with rapamycin, or starved for amino acids with and without ConA and used Ni-NTA beads to enrich ubiquitinated proteins. Anti-TNIP1 immunoblotting validated the MS findings and characterized TNIP1 as increasingly ubiquitinated in cells in which lysosomal degradation was inhibited (Fig 1D, suppl. Fig S1A).

To investigate if ubiquitination of Lys389 is necessary for lysosomal targeting of TNIP1, we performed site-directed mutagenesis and analyzed the stability and ubiquitination of respective TNIP1 variants. Next to Lys389 we also mutated the neighbouring residue Lys371, which we identified in two out of the three SILAC experiments. Arginine-variants did neither exhibit alterations in their global ubiquitination pattern, nor in their stability, indicating the presence of additional ubiquitination sites which are sufficient to support lysosomal degaradtion of TNIP1 (suppl. Fig S1B-D). Together, these results indicate that a multi-ubiquitinated form of TNIP1 accumulates upon the blockage of lysosomal acidification, and that TNIP1 may be an autophagy substrate. Due to its importance in inflammation, we decided to study the regulation of TNIP1 protein abundance in more detail.

### TNIP1 is degraded by autophagy

Because the blockage of lysosomal degradation by ConA led to an accumulation of ubiquitinated TNIP1, we examined whether TNIP1 is degraded by autophagy and if proteasomal degradation also contributes to regulating TNIP1 protein abundance under basal conditions. While inhibition of lysosomal acidification by ConA led to a significant accumulation of TNIP1 protein, inhibition of the proteasome by MG132 did not (Fig 2A). Furthermore, confocal immunofluorescent imaging showed endogenous TNIP1 accumulation upon Bafilomycin A1 (BafA1), another V-type ATPase inhibitor, treatment (suppl. Fig S2). Importantly under these conditions, TNIP1 colocalized with LC3-positive structures, which also strongly overlapped with p62/SQSTM1. Using airyscan super-resolution confocal microscopy, we found TNIP1 located inside LAMP1-and LC3-positive structures in untreated cells, and that TNIP1 accumulated within these structures upon BafA1 treatment (Fig 2B). Another way to test for lysosomal degradation is by using a tandem tag autophagy flux reporter system (Pankiv et al., 2007), in which TNIP1 is fused to a tandem mCherry-EYFP tag. While EYFP is sensitive to low pH, and therefore quickly loses its fluorescence in acidic lysosomes, the mCherry tag is rather stable under these conditions. In neutral cytosol, both tags of mCherry-EYFP-TNIP1 will be visible, while in lysosomes only the mCherry-tag will fluoresce and appear as a red-only dots. Transient transfection of mCherry-EYFP-TNIP1 did indeed lead to the formation of many red-only dots per cell, supporting the notion that TNIP1 ends up in lysosomal structures (Fig 2C). Taken together, this suggests that TNIP1 is degraded by autophagy under basal conditions as well as upon stress induced by rapamycin treatment and starvation, with proteasomal protein degradation playing a negligible role.

**Figure 2.**
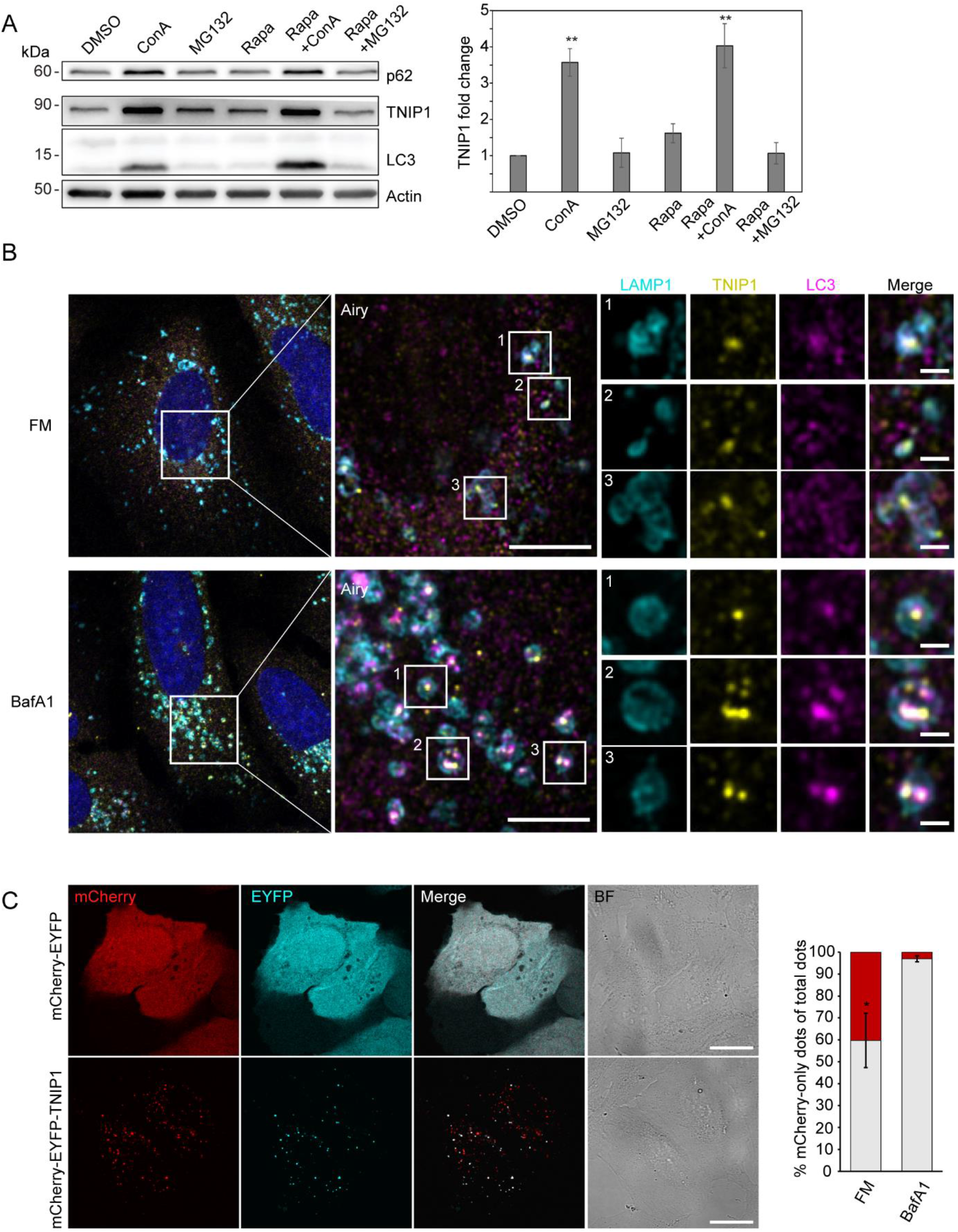
TNIP1 is degraded by autophagy. A U2OS cells were treated with 100 nM rapamycin for 4 h, proteasomal or lysosomal degradation were inhibited by 25 μM MG132 or 2 nM ConA, respectively. Under fed conditions (DMSO) and in rapamycin treated cells blockage of lysosomal acidification led to a significant increase of TNIP1 protein abundance (n=3). **: p<0.01 T test compared to DMSO treated samples. Error bars indiacte SEM. B Confocal images showing colocalization between endogenous TNIP1, LAMP1 and LC3 in U2OS cells treated for 5 hours with either BafA1 or vehicle DMSO. Cells were immuostained against endogenous TNIP1 (yellow), LAMP1 (cyan) and LC3 (purple) and imaged by Airyscan using the Zeiss LSM880 confocal microscope. Inserts highlight TNIP1 localized in LAMP1- and LC3-positive structures. Due to BafA1 treatment leading to accumulation of the immunostained proteins, signal intesities in the DMSO image have been increased relative to the BafA1 treated image during post-processing. Scale bars are 5 μm for the airyscan images, and 1 μm for the inserts. C U2OS cells were transiently transfected with either mCherry-EYFP or mCherry-EYFP-TNIP1. 24 hours after transfection, cells were either left untreated or treated with BafA1 for 4 hours. BF = bright field. Scale bars, 20 μm. Quantification of red-only TNIP1 dots over total TNIP1 dots was done using Volocity software (PerkinElmer), with intensity cut-offs based on BafA1 intensity of red and green dots (n=3). * = p<0.05, unpaired t-test. Error bars indicates SD.

Next, we aimed to characterize the molecular events that led to autophagosomal recruitment of TNIP1. We immunoprecipitated HA-tagged TNIP1 and performed a MS analysis of its interactome. This screen identified a number of autophagy-related proteins as possible TNIP1 interactors, including p62/SQSTM1, TAX1BP1, OPTN and TBK1 (Fig 3A-B, supplemental Table S2). The interaction with p62/SQSTM1 was also observed by immunoprecipitation followed by western blotting and appeared to be independent of autophagy activity, as neither starvation nor rapamycin treatment led to an increased interaction (Fig 3C). Staining of endogenous TNIP1 in U2OS cells further showed colocalization between TNIP1, p62/SQSTM1, TAX1BP1, and NDP52, other known SLRs, and that these accumulated together upon BafA1 treatment (Fig 3D). Thus, under basal and autophagy-inducing conditions TNIP1 gets ubiquitinated and interacts with SLRs, which likely target TNIP1 to autophagosomes for lysosomal degradation.

**Figure 3.**
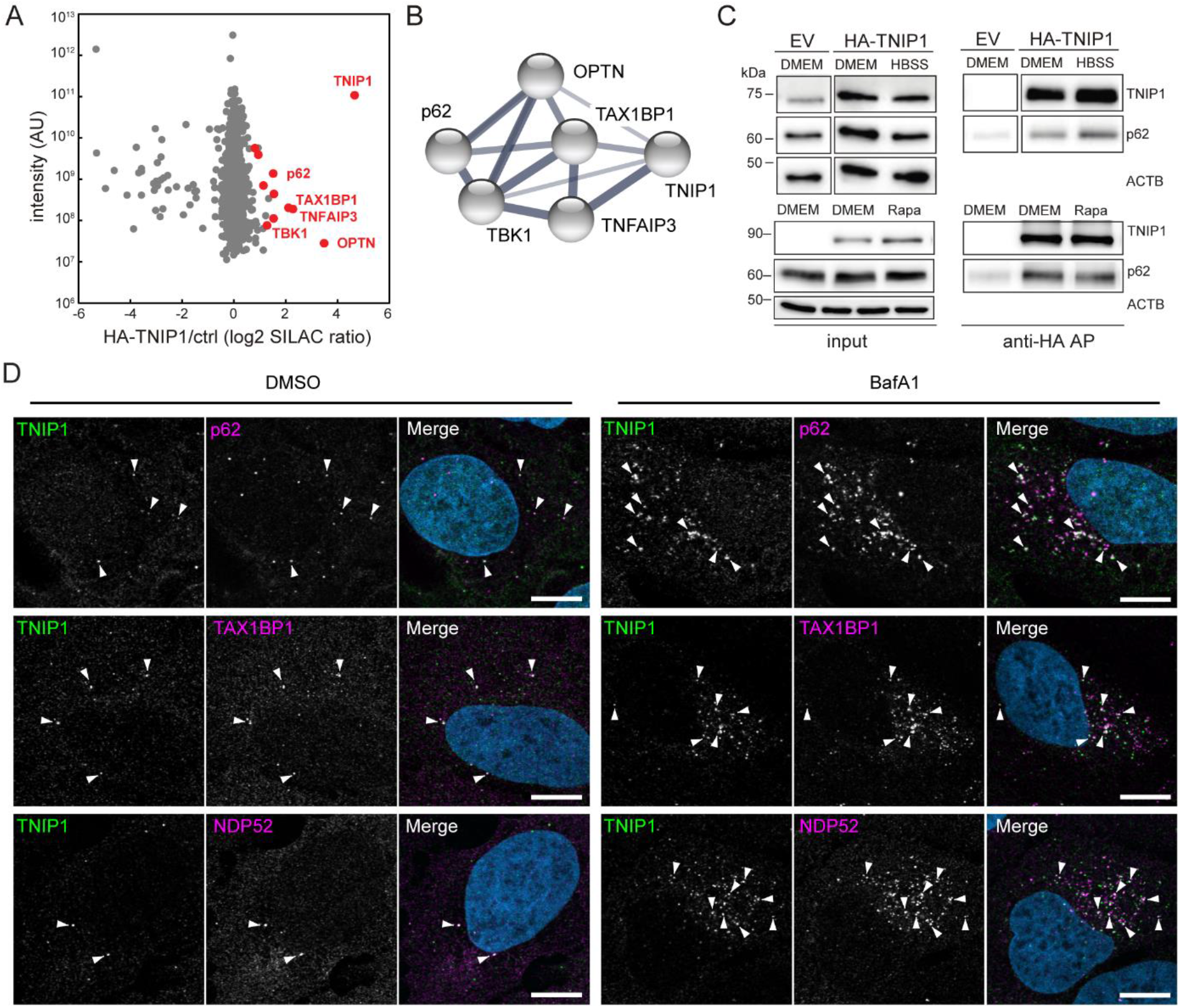
TNIP1 localizes to p62 bodies. A HA-TNIP1 affinity purification (AP)-MS highlights its interaction with autophagy receptors. HeLa cells expressing HA-TNIP1 and vector control cells (ctrl.) were differentially SILAC-labeled and anti-HA APs were performed under basal conditions, followed by quantitative MS analyses (n=3). Proteins which were significantly enriched in minimum two out of three replicates are highlighted in red (p<0.05, BH corrected). Proteins with known functions in autophagy and inflammation are annotated. B TNIP1 interactome. STRING DB was used to highlight the TNIP1 interactome identified in (A) (Szklarczyk et al., 2019). Thickness of edges indicate confidence of interaction. C TNIP1 interacts with p62/SQSTM1. Anti-HA affinity purifications followed by western blot analyses were performed to test for HA-TNIP1-p62/SQSTM1 interactions as identified in (A) in basal (DMEM), stress conditions (amino acid starvation, HBSS) and after rapamycin (Rapa) treatment each for 4 h. EV: empty vector. D U2OS cells were treated with either vehicle (DMSO) or BafA1 for 5 hours and stained for endogenous TNIP1 (green) together with either p62, NDP52 or TAX1BP1. Representative images are shown. Colocalization between TNIP1 and respective SLRs are indicated by arrowheads. Scale bars, 10 μm.

### TNIP1 interacts with human ATG8 family proteins via LIR motifs

Having established that TNIP1 colocalizes with several SLRs and is degraded by autophagy, we aimed to examine whether TNIP1 may also function independently of SLRs in autophagy. TNIP1 contains coiled coil domains for oligomerization and a ubiquitin-binding UBAN domain similar to that of OPTN, which preferentially binds K63-and M1-linked polyubiquitin chains (Fig 4A) (Herhaus et al., 2019; Wagner et al., 2008). TNIP1 only needs a LIR motif interacting with ATG8 proteins to be able to function as a SAR itself. Hence, we performed a peptide array screen for potential LIR motifs in human TNIP1. The peptide array, containing overlapping 20-mer peptides of TNIP1 moved by increments of three amino acids to cover the 636 amino acids full-length sequence, was probed with GST-GABARAP and revealed two potential LIRs in TNIP1, with core sequences 83-FDPL-86 and 125-FEVV-128, respectively (Fig 4B). Of these two candidates only LIR2 is conserved in the evolution of vertebrates down to cartilagous fishes, while LIR1 is only conserved down to marsupials and is not found in platypus (Fig 4C). LIR2 also has acidic- and phosphorylatable residues flanking the core LIR motif making it a very strong candidate for a functional LIR that could be positively regulated by phosphorylation (Johansen and Lamark, 2020; Wirth et al., 2019). LIR1 has a proline within the core motif which is usually inhibitory to binding to the ATG8s (Alemu et al., 2012; Johansen and Lamark, 2020). We then further tested the interaction between TNIP1 and the six human ATG8 family proteins by GST-pulldown assays. GST and GST-tagged human LC3 and GABARAP proteins were used to pull down *in vitro* translated wild-type TNIP1. Here, we observed that TNIP1 bound very well to several of the human ATG8s, with the strongest interaction being with LC3A, LC3B and GABARAP, and weak binding to LC3C and GABARAPL2 (Fig 4D-E). To test whether any of the two potential LIRs identified in the peptide array scan were responsible for this interaction, we mutated the conserved aromatic- and hydrophobic residues in each core LIR sequence to alanine, namely F83A/L86A and F125A/V128A (Fig 4C). GST-pulldown assays with these mutants revealed that the F125A/V128A mutations in LIR2 clearly reduced the interaction between TNIP1 and the GST-ATG8s, while the F83A/L86A mutations in LIR1 had a minor effect. Mutating both LIRs in TNIP1 further reduced the binding to some ATG8s, compared to mutating LIR2 alone (Fig 4D-E). We also tested the influence of LIRs for *in vivo* interactions of TNIP1 with LC3s using HeLa cells expressing HA-TNIP1 variants by performing anti-HA affinity purifications (APs) followed by anti-LC3A/B western blots (Fig 4F-G). In agreement with the *in vitro* observations, TNIP1 with both LIR motifs mutated (mLIR1+2) interacted significantly weaker with LC3A/B *in vivo*. Thus, TNIP1 fulfills all structural characteristics of being an SLR, with LIR2 being the motif mainly responsible for binding to ATG8s.

**Figure 4.**
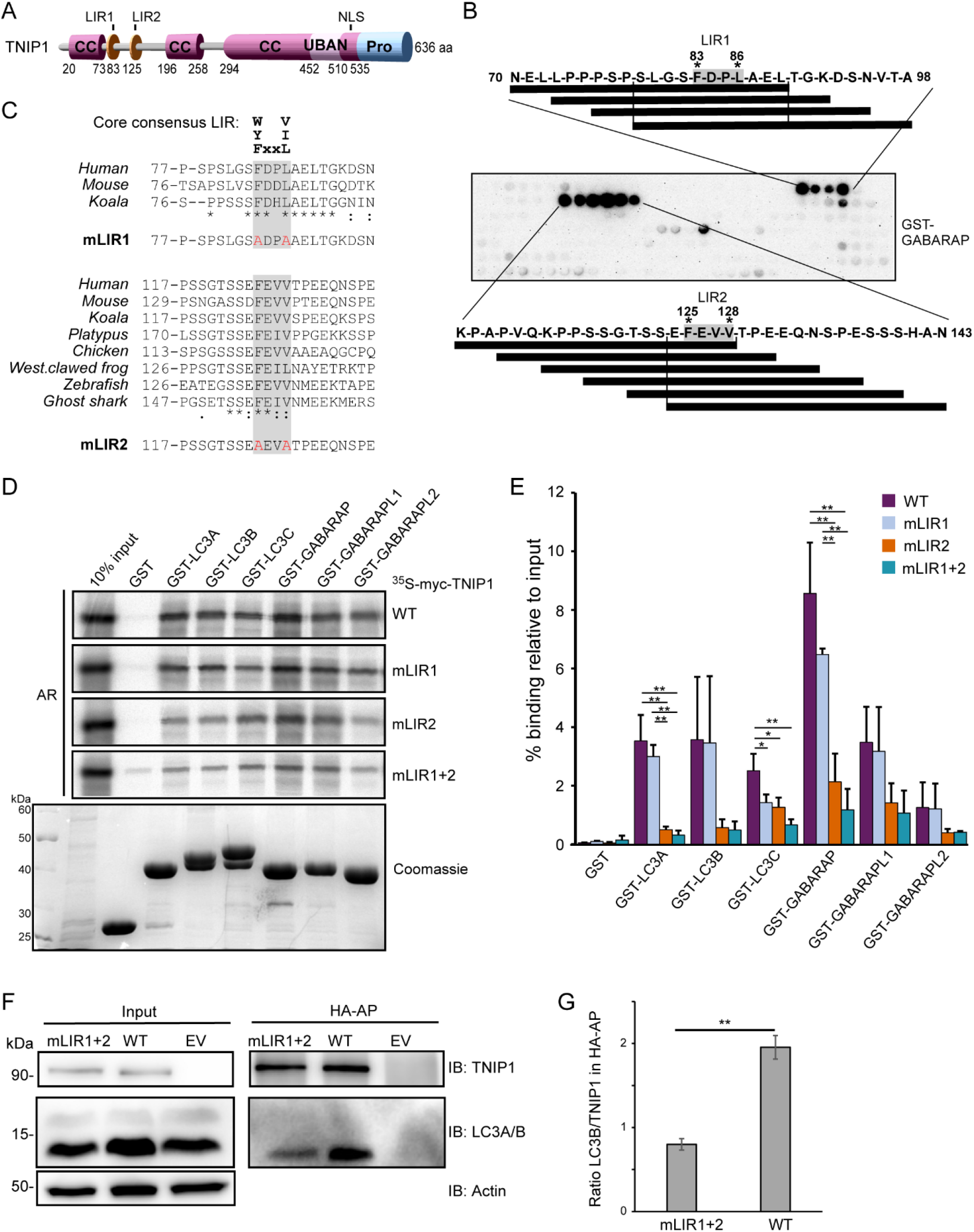
TNIP1 interacts with human ATG8 family proteins through a LIR motif. A Schematic drawing of the domain architecture of TNIP1, showing possible LIRs. B Peptide array of 20-mer peptides covering full length TNIP1 was used to probe for possible LIRs, using GST-GABARAP. C Amino-acid sequence alignment showing conservation of the core consenesus LIRs in TNIP1 across species. TNIP1 amino acid sequences were collected from UniProt, and multiple sequence alignment performed with Clustal Omega. Asterisk (*) indicates fully conserved residues. colon (:) indicates conservation between groups of strongly similar properties (>0.5 Gonnet PAM 250 matrix), and a period (.) indicates conservation between groups of weakly similar properties (0–0.5 Gonnet PAM 250 matrix). Mutated residues for the LIR mutants (mLIR1 and mLIR2) are shown in red. D *In vitro* GST-pulldown assay using 35S-labeled myc-TNIP1, myc-TNIP1-F83A/L86A (mLIR1), myc-TNIP1-F125A/V128A (mLIR2) and myc-TNIP1-F83A/L86A/F125A/V128A (mLIR1+2) against recombinant GST and GST-tagged human ATG8s. Bound myc-TNIP1 WT and LIR mutants were detected by autoradiography (AR). E Quantification of GST-pulldown from (D). Relative % binding was quantified against 10% input (n=3). * = p<0.05, **p<0.01, based on one-way ANOVA (post hoc: Tukey test). Error bars indicate SD. F LIR mutation impacts *in vivo* interaction with LC3A/B. Anti-HA AP of cells expressing HA-TNIP1^WT^ and HA-TNIP1^mLIR1+2^ were performed followed by western blot against indicated proteins. LIR mutation reduced the interaction between TNIP1 and LC3A/B. G Quantification of blots exemplified in panel (F) (n=3). Error bars indicate SEM. ** = p<0.01, T test.

### TNIP1 does not affect basal autophagy flux

We next asked whether TNIP1 was capable of regulating autophagy flux. For this we compared the levels of several known SLRs as well as the ratio of LC3-I and II under basal and starvation conditions in WT HeLa and two TNIP1 KO clones. The ratio of LC3-I and LC3-II was not affected by TNIP1 KO, and neither was the conversion upon starvation by HBSS, suggesting that TNIP1 does not affect autophagic flux under the tested conditions (Fig 5A-B). With the exception of OPTN, the basal protein levels of several SLRs and their degradation upon starvation was also unaffected by TNIP1 KO. However, the basal protein levels of OPTN were elevated in both TNIP1 KO clones, indicating that OPTN might be involved in a compensatory response. Indeed, we identified an upregulation of OPTN mRNA in TNIP1 KO clones (supplemental Table S3).

**Figure 5.**
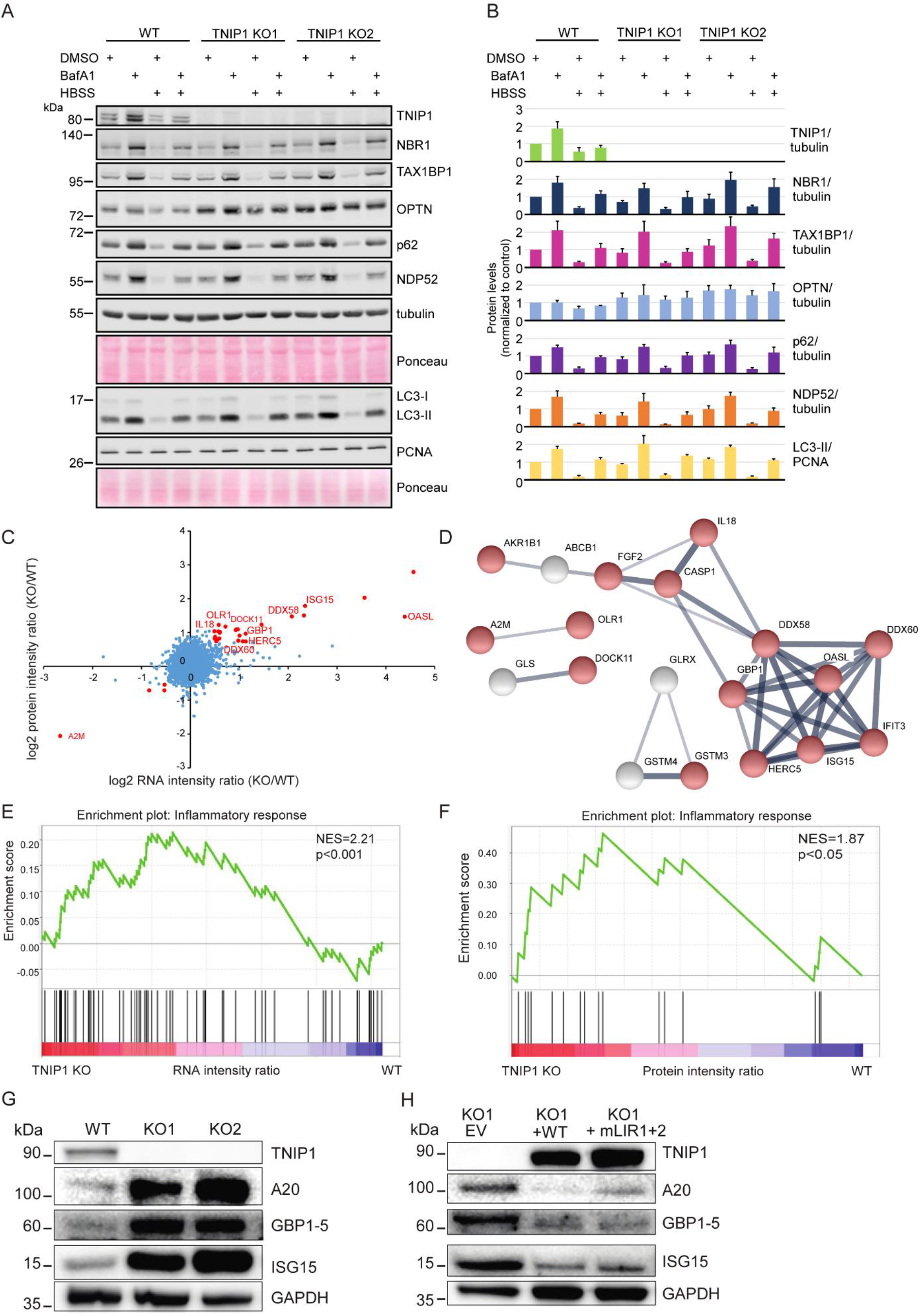
Loss of TNIP1 leads to an increase in inflammatory proteins. A Loss of TNIP1 does not alter autophagy flux under basal and starvation conditions. TNIP1 knock out HeLa cells generated by CRSPR/Cas9 were used to study its effect on autophagy. Wild-type HeLa cells and the two TNIP1 knockout clones denoted KO1 and KO2 were kept in either fed or starved (HBSS) conditions and treated with either vehicle (DMSO) or BafA1 for 8 hours. Blots were probed for several known SLRs as well as LC3. B Quantification of blots shown in panel (A) (n=3). Error bars indicates SD. C Loss of TNIP1 leads to an increased transcription of inflammatory genes. Fold changes of RNA and protein intensities of TNIP1 KO and WT cells were compared. Shown are average values of two KO clones compared to WT cells (n=3 per cell type for RNAseq; n=5 per cell type for proteomics). Genes that were significantly regulated on RNA and protein level are highlighted in red (p<0.01). Genes linked to immune effector processes and interferon-stimulated genes are annotated. D Protein-protein interactions of significantly regulated proteins. Proteins highlighted in red in (C) were analyzed on known interactions using STRING DB (Szklarczyk et al., 2019). Interactions between 17 proteins were identified, of which 13 are linked to stress response (marked in red). Thickness of edges indicate confidence of interaction. E, F Gene set enrichment analysis of significantly dysregulated mRNAs and proteins identifies an increased transcription and translation of genes involved in inflammation. NES denotes normalized enrichment score G, H TNIP1 represses translation of pro-inflammatory gene products. Whereas knockout of TNIP1 led to an increased abundance of indicated inflammatory proteins (G), re-expression of TNIP1^WT^ or TNIP1^mLIR1+2^ blunted this phenotype (H).

As SLRs are themselves autophagy substrates, and as loss of TNIP1 was shown to lead to increased inflammatory signaling (Shamilov and Aneskievich, 2018), we next asked if loss of TNIP1 in our cell systems also led to an increase of pro-inflammatory proteins. Comparing gene expression by RNAseq and protein abundance by SILAC-based proteomics between WT and TNIP1 KO HeLa cells, we indeed observed an upregulation of a number of genes involved in inflammatory signaling in TNIP1 KO cells on mRNA and protein level, including TNFAIP3, ISG15 and GBP1 (Fig 5C-D, supplemental Table S3). This was confirmed by gene set enrichment analyses (GSEA), which highlighted an activation of the inflammatory response in TNIP1 KO cells at both mRNA and protein level (Fig 5E-F). The upregulation of single proteins was also observed by western blot analysis (Fig 5G). Importantly, re-expression of TNIP1 blunted this effect confirming that it was indeed the absence of TNIP1 that led to the observed changes of the respective proteins. The LIR motifs did not affect this response under basal conditions as expression of TNIP1-mLIR1+2 led to the same consequences as TNIP1-WT (Fig 5H). Taken together, whereas TNIP1 seemed not to affect autophagy flux under basal conditions, its loss led to an increased abundance of pro-inflammatory proteins, indicating that selective autophagy may contribute to the tuning of inflammatory signaling by regulating TNIP1 protein levels. Thus, in this context TNIP1 is a *bona fide* autophagy substrate whose protein level is critical for the regulation of inflammatory signaling.

### Pro-inflammatory signaling induces LIR-dependent, autophagic degradation of TNIP1

So far, we characterized TNIP1 as a constitutive autophagic cargo as we did not observe changes in autophagosomal recruitment and lysosomal degradation based on the metabolic status of cells. To test if inflammatory signaling could lead to specific effects, we treated cells with the double-stranded RNA mimic poly(I:C), which mimics viral infection and elicits a TLR3 signaling response (Glavan and Pavelic, 2014). After 4 hours of poly(I:C) treatment we observed a significant decrease in endogenous TNIP1 levels, followed by an increase after 6 hours (Fig 6A-B). Parallel to the observed decrease of TNIP1, an increase in pro-inflammatory proteins like ISG15 was observed (suppl. Fig S3A). This decrease was dependent on canonical autophagy, as loss of ATG7 inhibited the response, but independent of SLRs, as loss of the SLRs p62/SQSTM1, NBR1, NDP52, TAX1BP1 and OPTN in a pentaKO cell line did not inhibit the response (Fig 6A-B). Hence, TNIP1 is itself specifically targeted to autophagosomes under poly(I:C) treatment. The higher TNIP1 levels seen in the pentaKO cells under basal conditions most likely indicate the involvement of the SLRs in basal turnover of TNIP1 (Fig 6A-B). Confocal imaging showed that upon poly(I:C) treatment in WT cells, endogenous TNIP1 changes from being mostly diffuse to forming dots in the cytoplasm (Fig 6C). The formation of TNIP1 dots in response to poly(I:C) was also observed in ATG7- and pentaKO cells, however, the phenotype of TNIP1 differed (Fig 6C). In ATG7 KO cells there appeared to be an increased amount of TNIP1 dots already at the basal level, which increased even further upon poly(I:C) treatment. Meanwhile, we observed an increased diffuse population in pentaKO cells at the basal level, most likely reflecting the increased protein levels observed in Fig 6A B. In pentaKO cells, TNIP1 responded similarly to poly(I:C) as WT cells, with a clear formation of dots. This suggests that SLRs are not necessary for poly(I:C)-induced TNIP1 aggregation into dots, and also not necessary for TNIP1 degradation upon poly(I:C) stimulation. It is possible that these dots represent the TNIP1 that is destined for degradation, and that the high amount observed in ATG7 KO cells is the result of blocked degradation.

**Figure 6.**
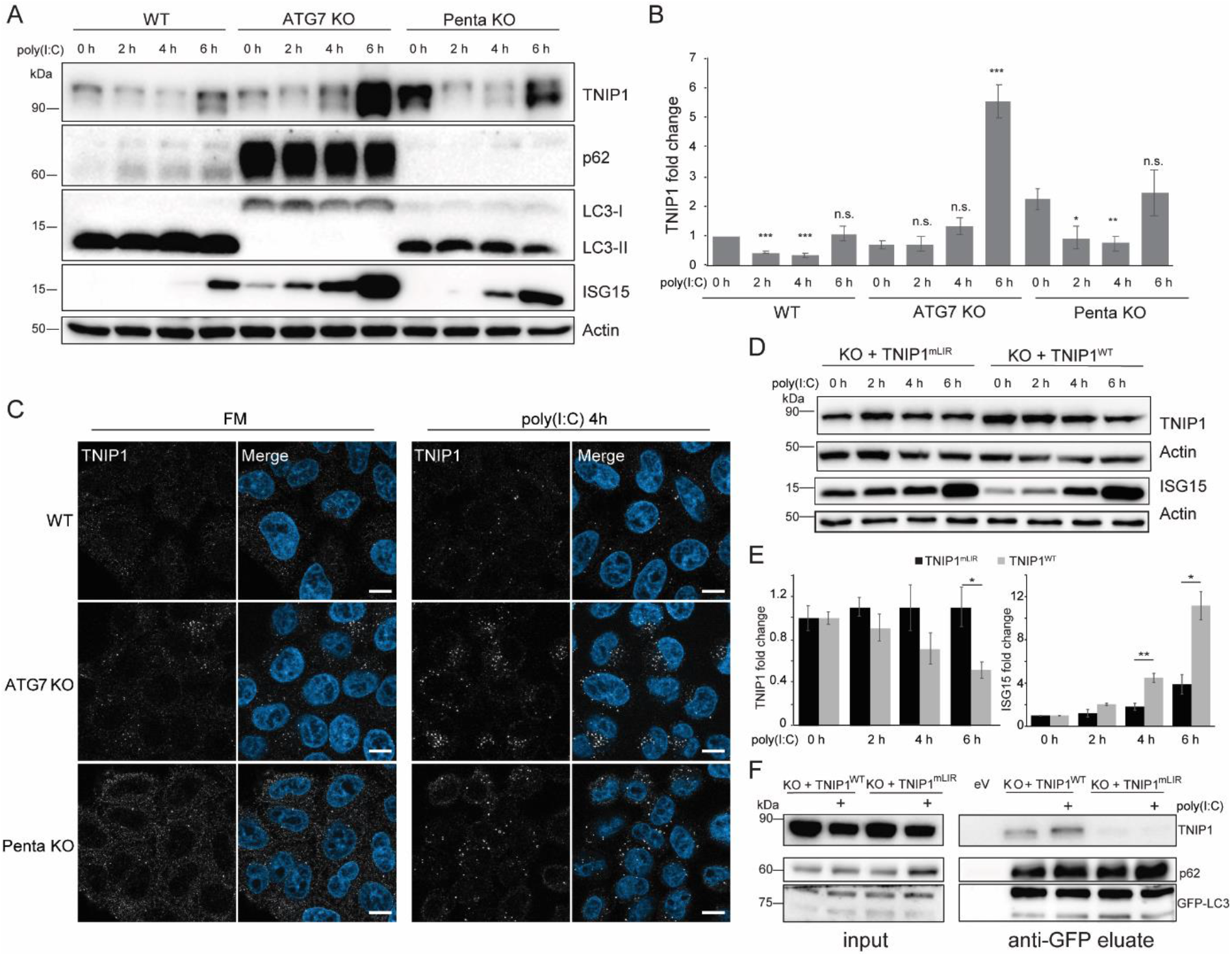
Poly(I:C) stimulation induces LIR-dependent, specific degradation of TNIP1 by autophagy. A Poly(I:C) treatment leads to time dependent changes in TNIP1 abundance. Poly(I:C) stimulation leads to an autophagy dependent decrease of TNIP1 abundance within the first 4 h as indicated by a block of degradation in ATG7 KO cells. Autophagy receptors appear to have a minor influence as degradation still occurs in pentaKO cells. B Quantification of blots shown in (A) (n=3). Error bars indicate standard deviation. * = p<0.05, ** = p<0.01, *** = p<0.001 unpaired T test compared to 0 h values of respective cell lines. C Representative immunofluorescent images showing endogenous TNIP1 response to poly(I:C) in WT, ATG7 KO and pentaKO. Cells were either left untreated or treated with 5 μg/ml poly(I:C) for 4 hours. Scale bar = 10 μm. D, E The degrdation of TNIP1 depends on functional LIR motifs. TNIP1WT is degraded in a time dependent fashion after poly(I:C) stimulation. The double LIR mutant TNIP1 (LIR1+2, TNIP1^mLIR^) is spared from degradation. Note: protein amounts of TNIP1 and ISG15 correlate inversely. Due to ectopic expression of TNIP1 variants regulation based on transcriptional/translational control as shown in panel (A) is lost. Panel (E) shows quantification of blots exemplified in panel (D) (n=3). Error bars indicate SEM. * = p<0.05, unpaired T test. F Poly(I:C) induces a LIR-dependent interaction with LC3. Indicated HeLa cells expressing GFP-LC3 were used for anti-GFP AP. Cells expressing TNIP1^mLIR^ do not exhibit an increased interaction between TNIP1 and GFP-LC3 after poly(I:C) treatment, in contrast to cells expressing TNIP1WT.

Because our data suggested that TNIP1 is degraded specifically by autophagy upon poly(I:C) exposure, we investigated whether this was LIR-dependent. To this end, we observed that in cells reconstituted with HA-TNIP1-WT, 6 hours of poly(I:C) treatment led to a significant decrease in TNIP1 levels, while we did not observe this decrease in cells expressing HA-TNIP1-mLIR1+2 (Fig 6D-E). ISG15 levels anticorrelated with HA-TNIP1 levels confirming the inhibitory role of TNIP1 in regulating ISG15 expression (Fig 6D-E). Importantly, the increase of ISG15 was significantly higher in HA-TNIP1-WT compared to HA-TNIP1-mLIR1+2 expressing cells, supporting the interpretation that LIR-dependent degradation of TNIP1 is critical for a stimulus- and time-dependent expression of pro-inflammatory genes. Note that the time-dependent increase of endogenous and exogenous TNIP1 after prolonged poly(I:C) treatment differed, indicating a long term transcriptional and/or translational regulation next to the observed short term autophagy-dependent effects (Fig 6D-E, suppl. Fig S3B). TNIP1 expression has previously been shown to be transcriptionally regulated by NF-kB, making it likely that prolonged poly(I:C) treatment can lead to the increased expression of TNIP1 (Tian et al., 2005). To further test the LIR-dependent recruitment of TNIP1 to autophagosomes under poly(I:C) treatment, we performed anti-GFP- LC3B IPs followed by anti-TNIP1 western blotting. Also in these IP experiments we observed a poly(I:C)-dependent increase in interaction of WT TNIP1 with LC3B in TNIP1 KO cells reconstituted with WT TNIP1, which was blocked in KO cells reconstituted with TNIP1- mLIR1+2 (Fig 6F). Thus, pro-inflammatory signling as exemplified by poly(I:C) treatment appears to lead to the specific, LIR-dependent degradation of TNIP1 by autophagy.

TBK1 positively regulates LIR-dependent interaction of OPTN with LC3B by phosphorylating the LIR motif of OPTN (Wild et al., 2011). In addition, TBK1 is an important mediator of TLR3-inducd antiviral signaling (Louis et al., 2018). Since we identified TBK1 as a possible interaction partner of TNIP1 (Fig 3A), we tested if TBK1 also plays a role in selective TNIP1 degradation. Indeed, confocal imaging showed that upon poly(I:C) treatment in WT cells, several of the observed endogenous TNIP1 dots colocalized with phosphorylated (Ser172), i.e. active, TBK1 (Fig 7A). Poly(I:C) treatment also led to a time-dependent activation of TBK1 as indicated by phosphorylation of Ser172 (Fig 7B). Interestingly, loss of TNIP1 led to an increased activation of TBK1 indicating a feedback regulatory mechanism.

**Figure 7.**
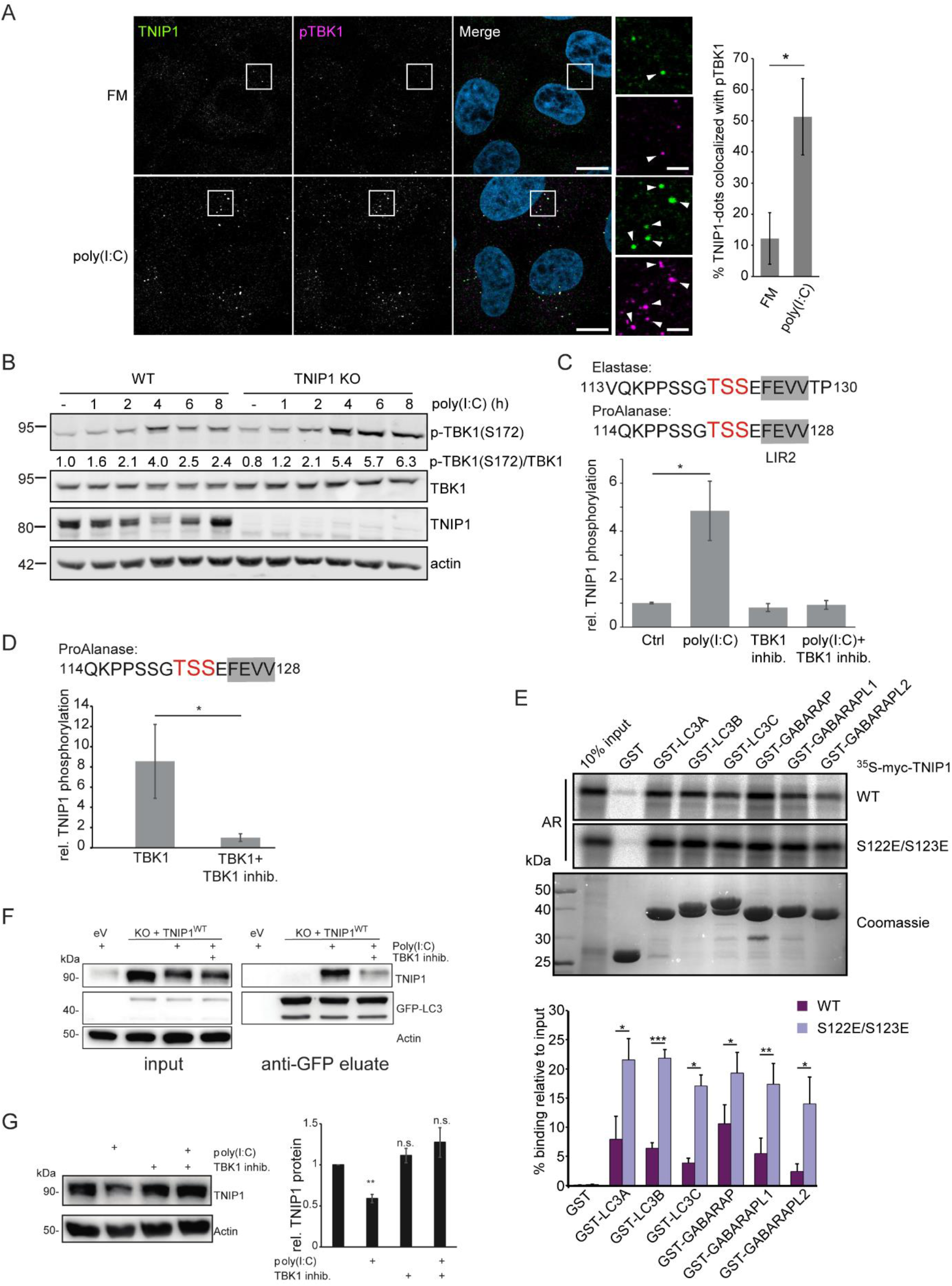
Poly(I:C) stimulation induces TBK1-dependent, specific degradation of TNIP1 by autophagy. A Immunofluorescence images showing colocalization between TNIP1 and pTBK1 upon poly(I:C) treatment. Cells were either left untreated or treated with 5 μg/ml poly(I:C) for 4 hours, and subsequently stained for endogenous TNIP1 and pTBK1. Colocalization between TNIP1 and pTBK1 is indicated by arrowheads. Quantification of TNIP1 dots colocalizing with pTBK1 was done using Volocity software (PerkinElmer). Around 160-220 cells were counted for each condition in each independent experiment (n=3). * = p<0.05, unpaired T test. Scale bar in overview image is 10 μm and scale bar in insert is 2 μm. B Time-course effect of poly(I:C) treatment on TBK1 activation and TNIP1. Representative blot and the corresponding quantification of the relative pTBK1 over total TBK1 levels are shown. C, D TBK1 phopshorylates TNIP1 N-terminal of LIR2. (C) *In vivo* phosphoproteomics using Elastase or ProAlanase as proteolytic enzymes identified indicated phosphopeptides. The single phosphorylation site could not be unambiguously localized to one of the three amino acid residues highlighted in red. Inhibition of TBK1 blocked the respective phosphorylation event (n≥3). Error bars indicate SEM. (D) *In vitro* kinase assay using purified TBK1 and TNIP1 coupled to phosphoproteomics indicates that TBK1 directly phosphorylates TNIP1 on one of the amino acid residues highlighted in red. E *In vitro* GST-pulldown assay using ^35^S-labeled myc-TNIP1 and myc-TNIP1-S122E/S123E against recombinant GST and GST-tagged human ATG8s. Bound myc-TNIP1 WT and S122E/S123E was detected using autoradiography (AR). n=3, * = p<0.05, ** = p<0.01, *** = p<0.001, unpaired t-test. Error bars indicate SD. F The interaction between TNIP1 and LC3B is regulated by TBK1. GFP-LC3B is purified using GFP trap beads. Bound TNIP1 is deteced by western blot. Inhibition of TBK1 by MRT67307 negatively regulates the poly(I:C)-dependent interaction of TNIP1 with LC3. G Inhibition of TBK1 negatively interferes with poly(I:C)-dependent degradation of TNIP1. Western blots of whole cell lysate indicate TNIP1 stabilization by TBK1 inhibition. Actin was used as loading control (n=3). Error bars indicate SEM. **: p<0.01, T test.

With the TBK1-mediated regulation of the OPTN LIR in mind, we used MS-based phosphoproteomics to study if TBK1 can phosphorylate serine and threonine residues located N-terminal to the core LIR sequence FEVV of LIR2 in TNIP1 (Fig 4C). Due to the surrounding amino acid sequence we could not follow the standard workflow of bottom-up proteomics experiments using trypsin to generate respective peptides. Instead, we performed a multi-protease digestion protocol using Elastase or ProAlanase to generate optimal sequence coverage (Eisenhardt et al., 2016; Samodova et al., 2020). We were able to identfy and quantify two phosphopeptide species which were phosphorylated within the TSS motif just in front of LIR2 (Fig 7C). Comparing cells treated or not with poly(I:C) and/or the TBK1 inhibitor MRT67307 clearly indicated that TBK1 phosphorylates TNIP1 in a stimulus-dependent manner (Fig 7C). To determine if TNIP1 is a direct target of TBK1 we performed *in vitro* kinase assays and could recapitulate the *in vivo* observations characterizing TNIP1 as a *bona fide* TBK1 substrate (Fig 7D). To test the effect of phosphorylations of the evolutionary conserved S122 and S123 residues N-terminal to the core LIR motif (Fig 4C), we analyzed the binding of the phosphomimicking S122E/S123E TNIP1 mutant to ATG8 family proteins by GST pulldown. Strikingly, the phosphomimicking mutant displayed strongly increased binding to all ATG8 proteins. The binding increase was particularly evident for LC3B, LC3C and GABARAPL2 (Fig 7E). Finally, we tested the effects of poly(I:C) treatment and TBK1 inhibition on TNIP1- LC3B interaction by GFP-LC3B IP. Inhibition of TBK1 led to a decreased interaction of TNIP1-WT with GFP-LC3B indicating that TBK1-dependent phosphorylation of the TNIP1 LIR motif was responsible for the observed increased interaction between LC3 and TNIP1 (Fig 7F). In agreement, inhibition of TBK1 led to a stabilization of TNIP1 under poly(I:C) treatment, i.e. interfered with its stimulus-dependent degradation (Fig 7G). Thus, whereas SLRs appear to support autophagy-dependent, basal turnover of ubiquitinated TNIP1, activation of its LIR motif by TBK1 induces its selective, autophagy-dependent removal, supporting the mounting of a transcriptional program to induce a robust inflammation response (Fig 8).

**Figure 8.**
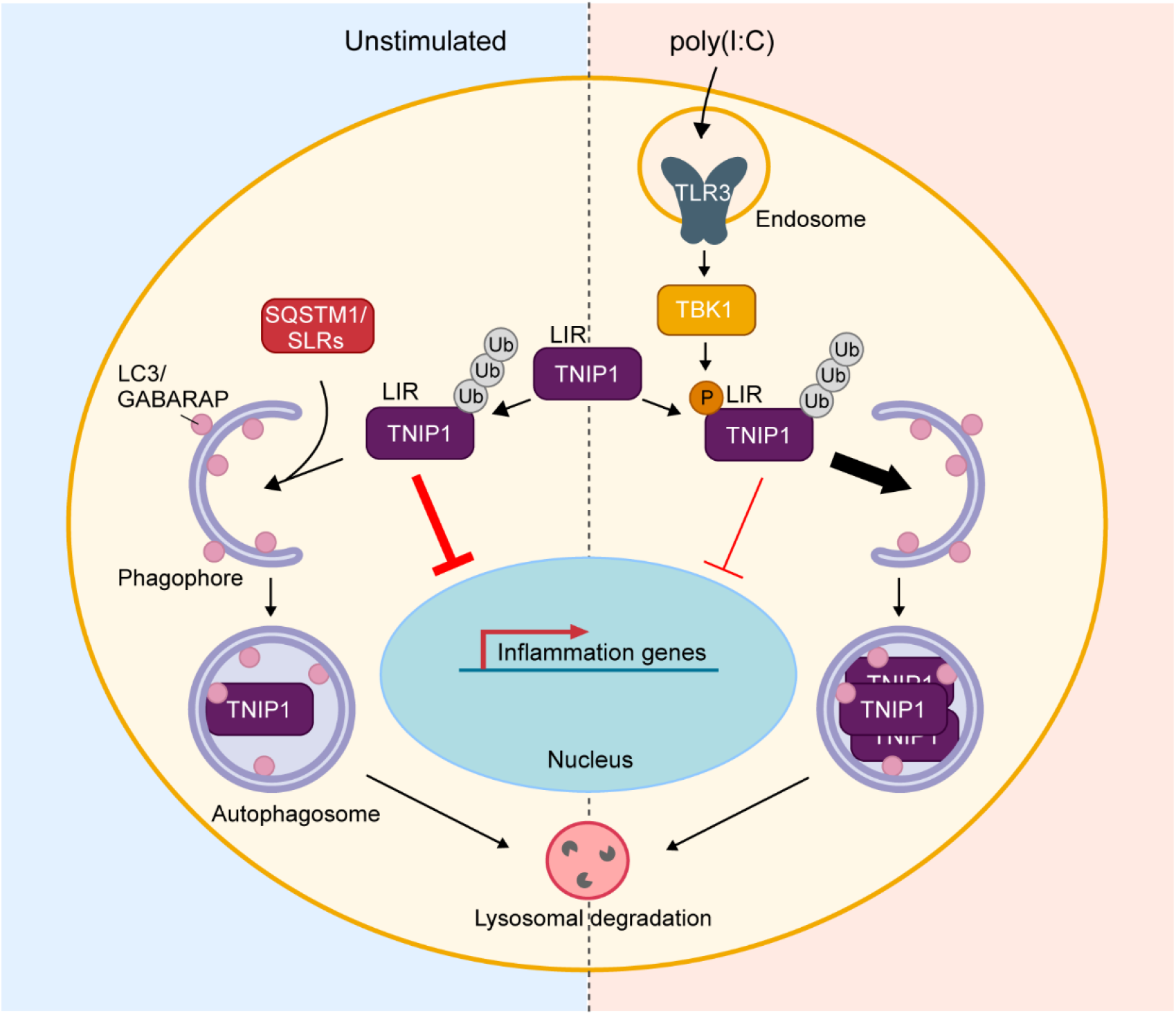
Model of TNIP1 regulation. Under basal, unstimulated conditions (left panel), TNIP1 functions as a negative regulator of inflammatory signaling, and is subject to constitutive autophagic degradation via interaction with autophagy receptors such as p62/SQSTM1. Upon poly(I:C)-induced activation of TLR3 (right panel), activated TBK1 phosphorylates TNIP1 in the vicinity of its LIR, increasing TNIP1 affinity for human LC3 and GABARAP proteins. This, in turn, leads to a LIR-dependent increase in TNIP1 degradation via selective autophagy. The removal of TNIP1 relieves the negative effect on inflammatory signaling, allowing the establishment of a robust inflammatory response upon antiviral signaling.

## Discussion

Autophagy is largely regarded as a cytoprotective, anti-inflammatory response ensuring cell and organismal homeostasis (Deretic, 2021; Deretic and Levine, 2018). Several autophagy loci have been linked to genetic predispositions for chronic inflammatory and autoimmune diseases. This indicates that lack of removal of damaged organelles may indirectly promote tissue destabilizing pro-inflammatory signaling. Autophagy may also contribute on a direct molecular level to limit excessive inflammatory signaling, e.g. by the selective degradation of inflammasome components by SARs, such as p62/SQSTM1 (Shi et al., 2012). Also, by removing other endogenous pro-inflammatory sources including damaged organelles and components from viral and bacterial infections, SARs play an important role in anti-inflammatory responses (Deretic, 2021). However, the pro-inflammatory functions of autophagy was so far limited to its contribution to interleukin secretion and to senescence-associated inflammation (Dupont et al., 2011; Lee et al., 2021; Zhang et al., 2015). In the current paper we highlight that the role of autophagy in inflammation appears to be more complex than anticipated and that both processes appear to be more intimately intertwined on a molecular level.

As TNIP1 has no reported enzymatic activity itself, it was believed that its functions in inflammation were directly linked to its interaction with TNFAIP3 (also known as A20), a ubiquitin-editing enzyme that contains both ubiquitin ligase and deubiquitinase activities and that was shown to negatively interfere with NFκ-B signaling (Shamilov and Aneskievich, 2018; Song et al., 1996). Several recent studies, however, indicate that TNIP1 itself may function as a key repressor of inflammatory signaling, its dysregulation being linked to hyperinflammatory diseases like psoriasis (Nair et al., 2009), systemic lupus erythematosus (Gateva et al., 2009), systemic sclerosis (Allanore et al., 2011), and senescence (Lee et al., 2021). TNIP1 is thus a potential target for the design of anti-inflammatory therapeutics. It has also been suggested that TNIP1 can function by out-competing other pro-inflammatory mediators for polyubiquitin binding, thereby negatively affecting inflammatory signaling (Shamilov and Aneskievich, 2018).

Whereas single nucleotide polymorphisms were shown to alter *TNIP1* expression and microRNAs to decrease *TNIP1* mRNA levels (Shamilov and Aneskievich, 2018), posttranslational mechanisms regulating TNIP1 protein abundance are largely unknown. We and others localized TNIP1 to autophagosomes, but did so far not address underlying mechanisms and phenotypical consequences (Mejlvang et al., 2018; Zellner et al., 2021). In the current study, we corroborate observations that TNIP1 can be regarded as a substrate for constitutive autophagosomal degradation. We did not identify changes in autophagy-dependent lysosomal degradation under fed and starved conditions, as well as under blocked MTORC1 signaling. The association of TNIP1 with SLRs and its colocalization with p62 bodies and LC3 puncta further supports this interpretation, indicating that ubiquitination of TNIP1 leads to SLR-dependent lysosomal degradation (Fig 8). Indeed, we could show that TNIP1 was still ubiquitinated even after mutating the two ubiquitination sites identified in this study. Thus, TNIP1 appears to be constitutively ubiquitinated and degraded by autophagy. This is in agreement to https://www.phosphosite.org which lists 11 ubiquitination sites of TNIP1, Lys389 being one of them (Akimov et al., 2018b).

Interestingly, this changes under inflammatory conditions. Poly(I:C) treatment led to SLR-independent, selective removal of TNIP1 by autophagy dependent on its LIR motif. This suggests the existence of two distinct TNIP1 pools within cells: one being phosphorylated and ubiquitinated and the other one being only ubiquitinated (Fig 8). This interpretation is supported by the observation that TNFAIP3 appeared to not be degraded by autophagy in a poly(I:C)- dependent manner, indicating that the pool of TNIP1 which interacts with TNFAIP3 under these conditions is also spared from degradation (see suppl. Fig S3A). The time-dependent regulation of TNIP1 protein levels indicates that autophagy-dependent degradation within the first 6 hours of poly(I:C) treatment contributes to establishment of a robust inflammatory response. After 6 hours TNIP1 levels rise again, most likely to prevent excessive inflammatory signaling which may lead to cell and tissue damage. In light of the model suggesting that TNIP1 can compete with pro-inflammatory proteins for ubiquitin binding (Shamilov and Aneskievich, 2018), it is possible that TNIP1 is degraded during the early stages of TLR3-activation to prevent this competition. The subsequent increase in TNIP1 at later stages of the signaling response could then be to outcompete other pro-inflammatory mediators, in order to limit excessive signaling. In a recent study OPTN was characterized as a direct, ubiquitination-independent binding partner of TNIP1 contributing to the specific, autophagy-dependent degradation of TNIP1 in senescent cells (Lee et al., 2021). The pathway characterized by us is different as TNIP1 degradation under inflammation conditions appears SLR independent (Fig 8). OPTN and TNIP1 appear to function in parallel and to be positively linked on the transcriptional level since TNIP1 KO led to an upregulation of OPTN mRNA and protein levels, potentially in a compensatory fashion.

The canonical LIR motif has the consensus sequence W/F/Y-X1-X2-L/I/V. The W/F/Y and the L/I/V occupy two hydrophobic pockets of the LIR docking site (LDS) of ATG8 family proteins. Additionally, acidic residues N-terminal, within and C-terminal, to the core motif, as found in the main LIR2 of TNIP1, increase binding affinity to the LDS which has a largely basic surface surrounding the two hydrophobic pockets (reviewed in Johansen and Lamark 2020). The presence of serine or threonine residues N-terminal to the core motif which can be phosphorylated to increase the acidic nature of the LIR is a neat strategy for a switchable LIR-LDS interaction. It has been shown for a number of SARs, including OPTN and several mitophagy receptors, that phosphorylation of N-terminal residues flanking the core LIR enhances the LIR-LDS interaction (Di Rita et al., 2018; Rogov et al., 2017; Wild et al., 2011; Wu et al., 2014; Zhu et al., 2013). This is also found for Beclin1, VPS34 and SCOC LIR-ATG8 interactions (Birgisdottir et al., 2019; Wirth et al., 2021). In the case of OPTN, the LIR-proximal phosphorylation is mediated by TBK1, leading to the enhanced binding of OPTN to LC3 (Wild et al., 2011). In this study we show that upon TLR3-activation, TNIP1 is phosphorylated at residues N-terminal to its main LIR-motif. Similar to OPTN, this enhances the interaction of TNIP1 with human LC3B. *In vitro*, phosphomimic S122E/S123E mutations of TNIP1 led to a strong increase in the binding to all of the human LC3 and GABARAP proteins, further supporting that the TNIP1 LIR is regulated by phosphorylation.

Furthermore, our results indicate that TBK1 is responsible for this phosphorylation. TBK1 is an important kinase in response to innate antiviral signaling. Upon TLR3-activation, TBK1 gets activated via TRIF and TRAF3 leading to the phosphorylation and activation of IRF3 and induction of type I interferons (Louis et al., 2018). TRIF itself may undergo autophagy-dependent degradation (Gentle et al., 2017; Inomata et al., 2012; Lim et al., 2019; Samie et al., 2018). How this anti-inflammatory effect of autophagy is coordinated with the here described pro-inflammatory acting degradation of TNIP1 will be an intersting question to address in future studies. TBK1 is also implicated in the regulation of autophagy, and is known to phosphorylate several autophagy proteins (Oakes et al., 2017). It is possible that TBK1 promotes TNIP1 degradation in order to ensure the efficient activation of downstream interferon signaling. Our results indicate that TNIP1 may rely on other SLRs for its basal turnover in unstimulated cells, but upon TBK1 activation in response to innate immune stimuli, the selective degradation of TNIP1 is promoted (Fig 8). Interestingly, TNIP1 fulfills all characteristics of a SAR, having a UBD, LIR motif and coiled-coil regions supporting multimerization. Whether TNIP1 mediates the degradation of selective cargo proteins under inflammatory conditions will have to be addressed in future studies. The knockout of TNIP1 resulted in a basal increase in the expression and protein levels of several pro-inflammatory proteins, even in the absence of pro-inflammatory signaling. This suggests that TNIP1 may have a role in preventing the induction of an inflammatory response even in untreated cells. Whether this is caused by TNIP1-mediated selective degradation of pro-inflammatory- mediators or −complexes, will be important to address in future studies. Taken together, we identified TBK1-dependent phosphorylation sites immediately N-terminal to the core LIR motif of TNIP1 that lead to its selective degradation by autophagy under inflammatory conditions. Autophagy may contribute to the establishment of a potent inflammatory response before it limits excessive cytotoxic inflammatory signaling. Thus in addition to cancer, inflammation is another condition in which autophagy may have a dual role either supporting or inhibiting underlying processes depending on the exact cell state and timing.

## Materials and Methods

The reagents, antibodies, and plasmids used in this study are listed in supplemental Tables S4, S5, S6, respectively.

### Cell culture and cell treatments

HeLa cells, and HEK293T cells were obtained from ATCC. U2OS cells were obtained from ECACC. StUbEx U2OS cells were a gift from Blagoy Blagoev (Department of Biochemistry and Molecular Biology, University of Southern Denmark, Odense, Denmark). HeLa CCL2.2, HeLa CCL2.2 ATG7 KO cells and HeLa CCL2.2 pentaKO cells were a gift from Richard J. Youle (National Institutes of Health, Bethesda, MD, USA). All cells were maintained in Dulbecco’s modified Eagle’s medium (DMEM) supplemented with 10% fetal calf serum, 2 mM L-glutamine, 100 U/ml penicillin/streptomycin. For starvation experiments, cells were incubated with Hanks’ Balanced Salt solution (HBSS) for the indicated times. Cells were treated with 100 nM rapamycin, 200 nM bafilomycin A1, 5 μg/ml Concanamycin A, 20 μg/ml cycloheximide or 10 μM MG132, for the indicated time periods. For SILAC, cells were grown for >14 days in SILAC DMEM supplemented with dialyzed fetal calf serum, 2 mM L-glutamine, 100 U/ml penicillin, and “light” (^12^C_6_^14^N_2_-Lysine and ^12^C_6_ ^14^N_4_-Arginine), “medium” (D_4_-Lysine and ^13^C_6_-Arginine), or “heavy” (^13^C_6_ ^15^N_2_-Lysine and ^13^C_6_ ^15^N_4_-Arginine) stable isotope-labeled amino acids. For transient DNA transfection, subconfluent cells were transfected using TransIT LT1 transfection reagent according to the manufacturer’s instructions.

### Plasmid constructs

Plasmids used in this study are listed in supplemental table S4. Cloning into pDest-vectors were done using the Gateway cloning system (Invitrogen). QuikChange site-directed mutagenesis kit (Stratagene) was used to create desired point mutations, which were verified by DNA sequencing (BigDye Sequencing kits, Applied Biosystems). Oligonucleotides for mutagenesis and sequencing were from Invitrogen. TNIP1 cDNA was obtained from Genscript (NM_001252390), TNIP1 LIR mutation fragment was synthesized by IDT gBlocks Fragments (Integrated DNA Technologies). TNIP1 and TNIP1_mLIR were cloned into pDONR201 and pLenti CMV Blast DEST (706-1) by Gateway recombination cloning according to manufacturer’s instructions. All plasmid constructs were verified by sequencing (BigDye; Applied Biosystems) and/or restriction digestion.

### Generation of knockout (KO) and stable cell lines

The TNIP1 KO cells were generated by CRISPR/Cas9-mediated genome editing tools provided by the F. Zhang laboratory (Broad Institute, MIT, Boston, MA). A target sequence in exon twenty of human TNIP1 was selected (sgTNIP1-1, 5’-CGTACCGGATCTACGACCCT-3’; sgTNIP1-2, 5’-GGCCCTGGAGTTCAACCGAC-3’; sgTNIP1-3, 5’-CACCCGACAGCGTGAGTACC-3’). The sgRNA oligos were cloned into the Bbs1 site of hSpCas9 plasmid. Lipofectamine LTX and Plus reagent (Invitrogen 15338100) were used to transfect hSpCas9-sgRNA into HeLa cells according to the manufacturer’s instructions. Cells were selected with 2 μg/ml puromycin for two days and single cells were isolated by serial dilutions. TNIP1 deficiency was screened by immunoblotting.

For generation of stable cell lines, lentiviral vectors were co-transfected with packaging (psPax2) and envelope plasmids (pMD2.G) in HEK293T cells by using jetPRIME reagent (Polyplus transfection). Medium was changed to fresh DMEM 15 h after transfection. After 24 h incubation, the supernatant was harvested by filtration through 0.22 μm filter. Polybrene (8 μg/ml) was added before infection of recipient cells. To obtain stable cell lines, infected cells were cultured in the presence of 5 μg/ml blasticidin for one week and monitored by immunoblotting.

### Enrichment of ubiquitinated peptides

Ubiquitinated peptides were purified according to Akimov et al. (Akimov et al., 2018a). Triple SILAC labeled U2OS and HeLa cells were treated with DMSO, Rapa, and Rapa+ConA. Cells were washed twice with cold PBS and lysed with 12 ml lysis buffer (8 M guanidine-HCl, 25 mM ammonium bicarbonate (ABC), pH 8.5). The lysates were sonicated to reduce viscosity and cleared by centrifugation at 15,000 RCF for 30 min. Protein concentration was determined by BCA protein assay kit (23225 and 23227 Pierce™) and the same amount of proteins of each label were mixed. Proteins were reduced with 2 mM DTT for 30 min at room temperature and alkylated with 11 mM CAA in dark for 30 min at room temperature. The concentration of guanidine-HCl was diluted to 2 M with 25 mM ammonium bicarbonate and filtrated with low binding 0.45 μm PVDF filters (Millipore). The Lys-C endopeptidase was added at a 1:100 enzyme to protein ratio to digest proteins at room temperature overnight. The peptide mixture was cleaned and purified using C18 cartridges (WATERS) and lyophilized for 24–36 h. The lyophilized peptides were dissolved in 12 ml of IAP buffer (50 mM MOPS pH 7.2, 10 mM sodium phosphate, 50 mM NaCl, PH 7.5-8.0) plus 0.1% Triton X-100. Dissolved peptides were spun down and passed through low binging 0.45 μm PVDF filters and incubated with 500 μl of UbiSite conjugated matrix for 5 h at 4 °C. The Ubisite matrix was washed 3 times with IAP buffer without detergent and 3 times with 150 mM NaCl. After 3^rd^ wash the matrix was transferred to a small Poly-Prep column (BioRAD). Enriched peptides were eluted with 250 μl 0.1% TFA 3 times, and each time incubated for 5 min. The eluted peptides were pooled and neutralized with 1 M ABC buffer to a final concentration at 25 mM, followed by trypsin digestion overnight at 37°C. The fractionation of tryptic peptides was performed as described previously (Akimov et al., 2018a). The resulting peptides were lyophilized and cleaned by STAGE-tips.

### Enrichment of ubiquitinated proteins by StUbEx

StUbEx U2OS cells were induced with doxycycline for 48 h and treated with Rapa and lysosomal degradation was blocked with ConA. The cells were washed with cold PBS and lysed with binding buffer (6 M Guanidinium-HCl, 50 mM Na2HPO4/NaH2PO4 pH 8.0, 500 mM NaCl, 5 mM imidazole). Sample viscosity was reduced by sonication followed by centrifugation at 11’000 RCF for 30 minutes at room temperature. Protein concentration was determined by BCA assay. Ubiquitin conjugates were purified by cOmplete His-tag purification beads (Roth) for 4 h at room temperature, followed by washes with binding buffer and washing buffers 1-3 (WB1 – 8 M Urea, 50 mM Na2HPO4/NaH2PO4 pH 8.0, 500 mM NaCl, 0.1% Triton-X-100; WB2 – same as WB1 but with 0.2% Triton-X-100; WB3 - 8 M Urea, 25 mM Tris pH 8.0, 150 mM NaCl). The ubiquitin conjugates were finally eluted with 3 times elution buffer (150 mM NaCl, 50 mM Tris pH 8.0, and 300 mM imidazole).

### Immunoblotting

For immunoblot analysis, treated cells were harvested and lysed in modified RIPA buffer supplemented with 2% SDS and 0.1% benzonase^®^ Nuclease or 1x SDS buffer (50 mM Tris pH 6.8, 2% SDS, and 10% glycerol). For cells harvested in modified RIPA buffer, lysates were centrifuged for 15 min at 13‘000 rpm. For cells harvested in 1x SDS buffer, lysates were boiled for 10 min. Protein concentration was measured using Pierce BCA Protein Assay Kit. Quantified cell lysates or AP elutes were either reduced with 2 mM DTT at 75 °C for 10 min or 100 mM DTT at 100°C for 10 min and resolved on SDS-PAGE gels. Proteins were transferred to a PVDF or nitrocellulose membrane, and subsequently prepared for either fluorescent or chemiluminescent detection. For fluorescent detection membranes were blocked in Intercept^®^ (PBS or TBS) Blocking Buffer, followed by overnight incubation at 4°C with primary antibody. Membranes were subsequently washed 4x in PBS or TBS containing 0.1% Tween (PBS-T/TBS-T), followed by incubation with secondary antibody diluted in Intercept^®^ Blocking Buffer. After 4x wash in PBS-T/TBS-T, a final wash was done in PBS/TBS without Tween. Fluorescent signal was detected using LiCOR Odyssey^®^ CLx imaging system. Chemiluminescent detection: membranes were blocked in either 5% milk or 5% bovine serum albumin in PBS-T or TBS-T, followed by incubation with primary antibody. Membranes were washed 3-4x in PBS-T/TBS-T followed by incubation with secondary antibody. After 3-4x washes in PBS-T/TBS-T, membranes were developed using SuperSignal West Pico Chemiluminescent Substrate on GE Fujifilm LAS4000 Luminescent image analyzer or Odyssey^®^ Fc reader (LI-COR Biosciences-GmbH). The densitometry of immunoblotting was performed by either ImageJ (National Institutes of Health) or ImageStudio software (LI-COR Biosciences-GmbH).

### Immunofluorescence staining and confocal fluorescence microscopy

For imaging, cells were grown on #1.5 round 12 mm coverslips (VWR, #631-0150) for immunofluorescence staining or eight-well Lab-Tek chamber coverglass for double-tag analysis (Thermo Fisher Scientific, #155411), and fixed with 4% formaldehyde in PBS for 30 min. For immunofluorescence staining, cells were washed 3x with PBS, and then permeabilized with either 0.1% Triton X-100 in PBS for 10 min, or ice-cold methanol for 10 min. Next, cells were washed 5x with PBS or TBS, and blocked for 1 h in 5% BSA in PBS or TBS. Subsequently, cells were incubated with primary antibodies diluted in 1% BSA in PBS or TBS for 1-2 h at room temperature. After 5x wash in PBS or TBS, cells were incubated with Alexa Fluor secondary antibodies diluted in 1% BSA in PBS or TBS for 1 h at room temperature. After 5x wash in PBS or TBS, cell nuclei were stained with 1 μg/ml DAPI diluted in PBS for 5 min, followed by 2x final washes in PBS or TBS. Coverslips were mounted using ProLong™ Gold or ProLong™ Glass Antifade Mountant. Cells were imaged using Zeiss LSM800 or LSM880 (Carl Zeiss Microscopy) using a 63× NA1.4 oil immersion lens for coverslips or a 40x NA1.2 water immersion lens for chambered coverglass. Images were collected in ZEN software (Zeiss). For Airyscan super-resolution images, optimal pixel size and z spacing as suggested by ZEN was used. Optimal excitation and emission settings were determined using the Smart Setup function. All fluorescence channels were recorded at non-saturating levels, and settings were kept identical between all samples within replicates used for comparisons or quantifications.

### Image analysis

For quantification of red-only dots of mCherry-EYFP-TNIP1, transiently transfected cells with low expression of mCherry-EYFP-TNIP1 were visually selected and imaged. Cells with high levels of overexpression were excluded, as this resulted in the formation of large aggregates. mCherry and EYFP dot detection was performed using a custom-made measurement protocol using intensity thresholding, size exclusion, and noise filtering, based on signal intensity of the BafA1 control in Volocity software (PerkinElmer) ver. 6.3. The number of mCherry-only positive dots was counted by subtracting total EYFP dots from total mCherry dots for each experiment. Around 2000-3000 dots were counted for each condition within each replicate.

To quantify the colocalization between TNIP1 and pTBK1, populations of objects representing fluorescent puncta in each channel were segmented using a custom-made protocol in Volocity ver. 6.3 (PerkinElmer). Detection of TNIP1 and pTBK1 puncta was performed by intensity thresholding, size exclusion and noise reduction. Overlap between TNIP1 and pTBK1 was identified by excluding TNIP1 objects not touching pTBK1. The percentage of TNIP1 dots colocalizing with pTBK1 was calculated by dividing the number of TNIP1 dots overlapping with pTBK1 with the total number of TNIP1 dots. Around 160-220 cells were counted for each condition within each replicate. For line-profiles, the line-profile tool in Zen Blue software (Zeiss) was used to measure signal intensity across indicated lines.

### SPOT synthesis and peptide array

TNIP1 peptide array was synthesized on cellulose membranes using MultiPrep peptide synthesizer (INTAVIS Bioanalytical Instruments AG, Germany). Membranes were blocked with 5% non-fat milk in TBS-T and peptide interactions were tested using GST-GABARAP by overlaying the membrane with 1 μg/ml recombinant protein for 2 h at room temperature. Membranes were washed three times in TBS-T. Bound GST-GABARAP was visualized with HRP-conjugated anti-GST antibody (Johansen et al., 2017). Putative LIR motifs in 20, 3 arrays (20-mer peptides moved a window of 3 residues along the protein sequence) were identified as 4–6 consecutive strong spots containing the core LIR consensus (W/F/Y)XX(L/I/V).

### Recombinant protein production and GST-pulldown analysis

GST and GST-fusion proteins were expressed in *Escherichia coli* strain SoluBL21 (DE3) (Genlantis, #C700200). Protein expression was induced by adding 50 g/ml isopropyl β-D-1- thiogalactopyranoside (IPTG). The bacterial cells were sonicated in lysis buffer (20 mM Tris-HCl pH 7.5, 10 mM EDTA, 5 mM EGTA, 150 mM NaCl) and GST-fused proteins were immobilized on Glutathione Sepharose 4 Fast Flow beads (GE Healthcare, #17-5132-01) by incubating in a rotator at 4°C for 1 h. The beads containing GST-fusion proteins were subsequently used for pulling down *in vitro* translated proteins or cell lysates. For *in vitro* translated proteins, a pDest-myc-vector containing the protein of interest and a T7 promoter was used. *In vitro* translation was done using the TNT T7 Reticolucyte Lysate System (Promega Corp.), in the presence of radioactive ^35^S-methionine. *In vitro* translated protein was then precleared by incubation with empty Glutathione Sepharose beads in 100 μl of NETN buffer (50 mM Tris pH 8.0, 150 mM NaCl, 1 mM EDTA, 0.5% NP‐40) supplemented with cOmplete Mini EDTA‐free protease inhibitor (Merck) for 30 min at 4°C. Precleared lysates were then incubated with GST-fusion protein bound beads for 1-2 hours on a rotator at 4 °C. Beads were then washed 5x with NETN-buffer, and resuspended in 2× SDS-PAGE gel loading buffer (125 mM Tris, pH 7.5, 4% SDS, 0.04% bromophenol blue, 20% glycerol, 100 mM dithiothreitol), boiled for 10 min, and resolved by SDS-PAGE. Gels were stained with Coomassie Brilliant Blue for protein visualization, and then vacuum-dried. The radioactive signal was then detected on imaging plates with Fujifilm BAS‐5000 (Fujifilm). Signals from ^35^S-labelled proteins were measured in terms of unit of photostimulated luminescent (PSL) and quantified in comparison with 10% of the *in vitro* translated lysate (input) using the Image Gauge software (Fuji) (Johansen et al., 2017).

### Affinity purification

Cells were lysed in ice-cold modified RIPA buffer (25 mM Tris-HCl pH 7.4, 150 mM NaCl, 1 mM EDTA, 1% NP-40, 0.1% Sodium deoxycholate) containing complete protease inhibitor cocktail and phosphatase inhibitor cocktail. Lysates were centrifugated for 15 min at 13’000 RCF and protein concentration was determined by BCA assay. GFP-tagged proteins were affinity purified by GFP-trap (ChromoTek) according to the manufacturer’s instructions. HA- TNIP1 was affinity purified by anti-HA magnetic beads (Pierce). TNIP1 polyclonal antibody coupled protein G dynabeads were used for affinity purification of endogenous TNIP1. For denaturing purification, the lysate was supplemented with 2% SDS and incubated for 30 min at room temperature to break protein-protein-interaction, then diluted to 0.5% SDS with lysis buffer and incubated with HA or anti-TNIP1 antibody for affinity purification.

For TNIP1 interactome analysis by MS, HA-TNIP1 cells and empty vector cells were cultured in “heavy” and “light” SILAC medium respectively. After two weeks, cells were harvested and lysed in modified RIPA buffer containing complete protease inhibitor cocktail and phosphatase inhibitor cocktail. Protein concentration was determined by BCA assay and protein amount was adjusted to equal concentration with lysis buffer, followed by affinity purification with anti-HA magnetic beads. The eluate of “heavy” and “light” samples was combined and fractionated by SDS-PAGE. Proteins were in-gel digested by trypsin and peptides were desalted by STAGE-tips prior LC-MS/MS analysis.

### Whole proteome analysis

For whole proteome analysis by MS, HeLa wild-type cells and TNIP1 KO cells were cultured in SILAC medium for two weeks. Cells were harvested and lysed in modified RIPA buffer containing 2% SDS and 0.1% benzonase^®^ Nuclease. Protein concentration was determined by BCA assay and an equal amount of protein in each labeling was mixed. Protein was reduced with DTT and alkylated with IAA, followed with SDS-PAGE fractionation and trypsin in-gel digestion. Tryptic peptides were desalted by STAGE-tips prior LC-MS/MS analysis.

### TNIP1 phosphorylation analysis

HA-TNIP1 cells were treated with poly(I:C) with or without TBK1 inhibitor MRT67307. Cells were harvested and lysed in modified RIPA buffer containing complete protease inhibitor cocktail and phosphatase inhibitor cocktail, 2% SDS was supplemented to break protein-protein interaction. The samples were diluted to 0.5% SDS with lysis buffer and subjected to affinity purification with anti-HA magnetic beads. After purification, the beads were transferred onto 10 kDa MW cut-off filter with 400 μl 8 M urea and 1 mM DTT. Proteins were digested by elastase or ProAlanase using the FASP protocol (Wisniewski et al., 2009). Digested peptides were eluted twice with 200 μl 50 mM ammonium bicarbonate into fresh tubes and acidified with TFA to a final concentration of 1%. Peptides were frozen and lyophilized prior to phosphopeptide enrichment. The lyophilized peptides were resuspended in 200 μl of 80% ACN, 0.1% TFA. Phosphopeptides enrichment was performed on Agilent AssayMAP Bravo platform (Basel, Switzerland) according to manufacturer’s instructions. The Agilent AssayMAP phosphopeptide enrichment v2.0 App was used for automated phosphopeptide enrichment using Fe (III)-NTA cartridges (Basel, Switzerland). Cartridges were primed with 100 μl of 50% ACN, 0.1% TFA using a high flow rate of 300 μl/min and equilibrated using 80% ACN containing 0.1% TFA. Samples were loaded onto the cartridge using a low flow rate of 3 μl/min. The cartridges were washed twice with 200 μl 80% ACN containing 0.1% TFA and eluted with 50 μl of 1% ammonium hydroxide (pH 11) and 50 ul of 1% ammonium hydroxide in 80% ACN to a low binding PCR tube containing 5 μl FA. The eluted peptides were lyophilized and resuspended in 20 μL of 0.1% FA for LC-MS/MS analysis.

### In *vitro* kinase assay

HA-TNIP1 cells were harvested and lysed in ice-cold modified RIPA buffer containing complete protease inhibitor cocktail. TNIP1 protein was enriched by anti-HA magnetic beads and dephosphorylated by lambda phosphatase with 1 mM MnCl2 for 30 min at 30 °C. Beads were washed with 2 x 10 mL of kinase buffer (50 mM Tris-HCl pH 7.6, 10 mM MgCl2, 150 mM NaCl and 1x PhosSTOP) to remove lambda phosphatase. Kinase buffer was added to a final volume of 800 μl with 1 μg TBK1 protein, for control sample, 10 μM TBK1 inhibitor MRT67307 was added. Kinase assays were performed on a rotor at 37°C for 1 h. Finally, reactions were quenched by addition of 8 M urea and 1 mM DTT. The beads were transferred onto 10 kDa MW cut-off filter for the digestion with ProAlanase, followed with phosphopeptide enrichment.

### LC-MS/MS analyses

Solubilized peptides were injected into a 20 cm fused silica column with an inner diameter of 75 μm and in-house packed with C18 (ReproSil-Pur 120 C18-AQ, 1.9 μm, Dr. Maisch) for reverse phase fractionation by EasyLC 1200 nanoflow-HPLC system (Thermo Fisher Scientific). Peptides were loaded with solvent A (0.1% FA in water) at a max. pressure of 800 Bar and eluted with a step gradient of solvent B (0.1% FA in 80% ACN) from 2% to 25% within 85 min, from 25% to 60% within 5 min, followed by increasing to 100% in 2 min at a flow rate of 250 nl/min. MS/MS analysis was performed on a nano-electrospray ion source equipped QExactive HF-X mass spectrometer (Thermo Fisher Scientific). The spray voltage was set to 2.3 kV with a capillary temperature of 250°C. Mass spectrometer was operated in positive polarity mode and MS data were acquired in the data-dependent mode. The automatic gain control (AGC) target was set to 3×10^6^, the resolution was set to 120’000, and the ion injection time was set to 15 ms for the full scan at a mass range of m/z=370 to 1750. Twelve precursors were fragmented using normalized collisional energy (NCE) of 28 by higher-energy collisional dissociation (HCD). MS/MS scans were acquired with a resolution of 30’000, AGC target of 100′000 maximum IT of 54 ms, isolation window of 1.6 m/z, and dynamic exclusion window of 30 s. MS raw files were analyzed using MaxQuant software version 1.6.2.10 (Cox and Mann, 2008), data were searched against UniProt full-length homo sapiens database (21.033 entries, released March, 2016). Cysteine carbamidomethylation was set as a fixed modification, protein aminoterminal acetylation and methionine oxidation were set as variable modifications. Ubiquitination of lysine was set as a variable modification and digestion was specific to trypsin/P for Ubisite experiment. Serine-, threonine- and tyrosine-phosphorylation were set as a variable modification and digestion was set to unspecific or ProAlanase for TNIP1 phosphorylation experiments. The analysis was carried out with “match-between-run” with a time window of 0.7 min. MaxQuant results were analyzed using Perseus (Tyanova et al., 2016).

### Total RNA extraction and library construction

Four replicates of HeLa WT cells and TNIP1 KO cells (clone1 and clone2) were cultured in 10 cm dishes. The total RNA was extracted using RNeasy Mini Kit (Qiagen) according to the instructions of the manufacturer. Cells were lysed with 350 μl RLT buffer and homogenized by passing 5 times through a blunt 20-gauge needle. One volume of 70% ethanol was supplemented, the lysates were transferred to RNeasy spin column and centrifuged at 8‘000 x g to collect RNA. The RNA was washed once with 700 μl RW1 buffer and twice with 500 μl RPE buffer. After completely removing the RPE buffer, the RNA was eluted with 50 μl RNAse- free water. The quality of RNA samples was analyzed by RNA screen tape (Agilent). The complementary DNA (cDNA) libraries were barcoded using Illumina primers and sequenced on one lane of an Illumina Novaseq 6000 instrument with 2×50 bp paired-end sequencing cycles. The sequence data were deposited at the European Nucleotide Archive (accession number: PRJEB45902). The output reads from multiple lanes were combined into single files and quality control was performed with FastQC v0.11.7 and cleaned with fastp v0.19.5 to removed polyG trails and keep only full-length reads (Chen et al., 2018). The human genome GRCh38.p13 (ENSEMBL) was used to remap the reads using STAR v2.5.3a (Dobin et al., 2013). The differential expression between WT and KO was analyzed by R software (R Core Team, 2014 http://www.R-project.org/) using the DESeq2 package (Love et al., 2014).

### Gene set enrichment analyses

Gene set enrichment analyses (GSEA) was performed by using default setting weighted enrichment statics and signal2noise metric for ranking genes (Subramanian et al., 2005). Significantly regulated genes in transcriptome data and proteins in whole proteome data were listed in supplemental table S3.

### Quantification and statistical analyses

Significantly regulated ubiquitination sites were determined by two samples paired t-test, FDR<0.05, S0=0.1. The interactome network was generated by the String database. The statistical analyses including standard deviations, error bars and P-values were performed using Excel (Microsoft), all the details were indicated in figure legends.

## Data availability

The mass spectrometry proteomics data have been deposited to the ProteomeXchange Consortium via the PRIDE (Perez-Riverol et al., 2019) partner repository with the dataset identifier PXD027163, Username, Password: nsna7vnu.

The RNA sequence data were deposited at the European Nucleotide Archive (accession number: PRJEB45902).

## Supplemental Material

- **Supplemental Figure S1.** TNIP1 gets ubiquitinated and degraded in the lysosome.
- **Supplemental Figure S2.** TNIP1 localizes to autophagosomes.
- **Supplemental Figure S3.** Regulation of TNIP1 protein abundance.
- **Supplemental Table S1.** UbiSite-proteomics approach to identify ubiquitination sites on proteins being potentially involved in autophagy.
- **Supplemental Table S2.** HA-TNIP1 interactome.
- **Supplemental Table S3.** RNA-Proteome correalation comparing WT and TNIP1 KO HeLa clones.
- **Supplemental Table S4.** Chemical reagents used in this study.
- **Supplemental Table S5.** Antibodies used in this study.
- **Supplemental Table S6.** Plasmids used in this study.

## Acknowledgements

The technical assistance of Aud Øvervatn is greatly appreciated. We thank Laurent Falquet for RNAseq data analysis. We are grateful to Richard J. Youle for the generous gift of the pentaKO cells. We thank the bioimaging core facility, Department of Medical Biology, UiT – The Arctic University of Norway for expert assistance. This work was funded by grants from: FRIBIOMED (grant 214448) and TOPPFORSK (grant 249884) programs of the Research Council of Norway to T.J.; the Swiss National Science Foundation, the Canton and University of Fribourg, and the Novartis Foundation for Medical-Biological Research to J.D. This work was part of the SKINTEGRITY.CH collaborative research project.

## Author contributions

JD, TJ, NLR and JZ designed the experiments. JZ and NLR performed most of the experiments. GE performed GST pulldown experiments and analysed data. VA and BB performed and analysed Ubi-Site experiments. ZH performed phosphoproteomics. HLO assisted in supervision performed *in vitro* mutagenesis and analysed data. TL analyzed data and assisted in supervision. TJ and JD supervised and co-ordinated the overall research, analyzed the data, and approved the final manuscript. JZ, NLR, HLO, TJ and JD wrote the manuscript.

