## Supplemental Figures for "TBK1 phosphorylation activates LIR-dependent degradation of the inflammation repressor TNIP1"

*****Should be considered equal first authors

### Supplemental Figures

- **Supplemental Figure S1. TNIP1 gets ubiquitinated and degraded in the lysosome.**
- **Supplemental Figure S2. TNIP1 localizes to autophagosomes.**
- **Supplemental Figure S3. Regulation of TNIP1 protein abundance.**

### Supplemental Figure S1

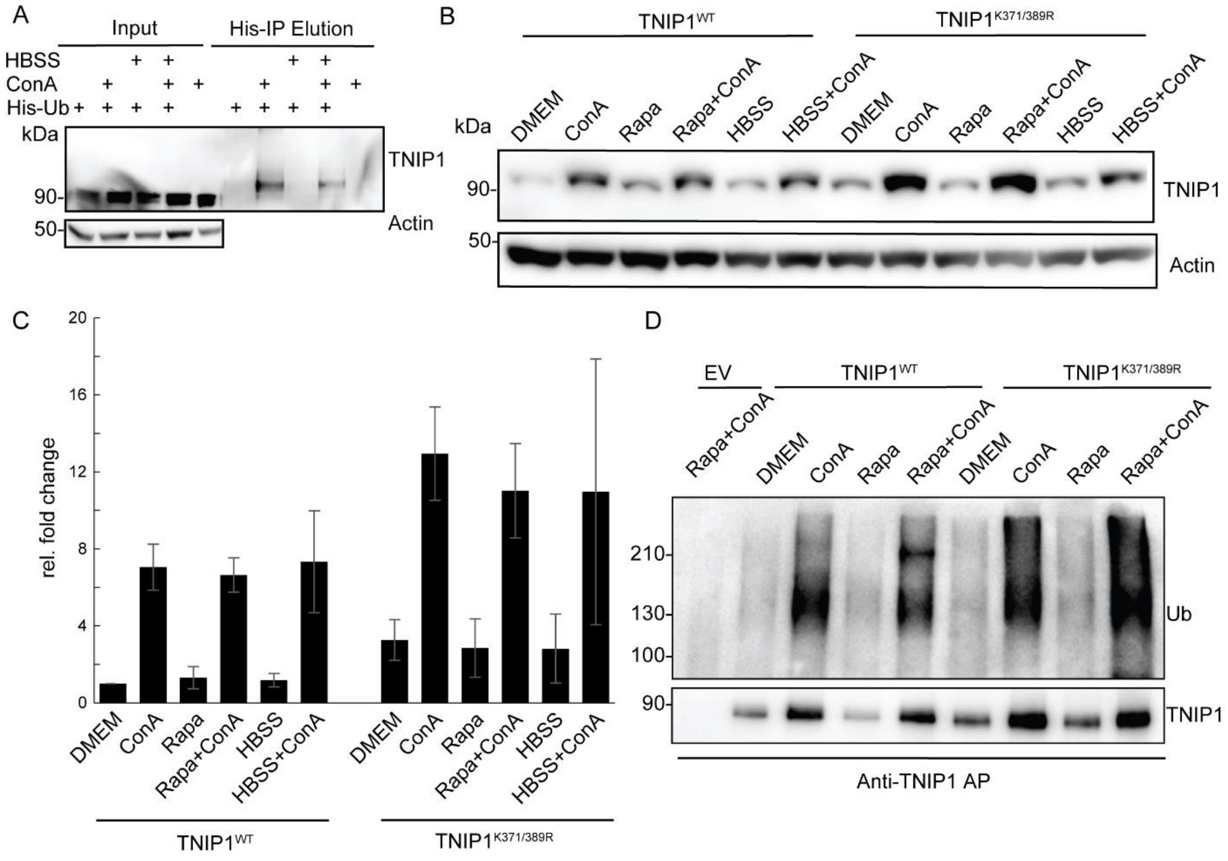


**Supplemental Figure S1. TNIP1 gets ubiquitinated and degraded in the lysosome.**

A U2-OS-StUbEx cells inducibly expressing His-FLAG-tagged ubiquitin at endogenous levels were used to enrich ubiquitinated proteins (Akimov et al., 2014). Under control conditions as well as under starvation treatment (HBSS) TNIP1 gets ubiquitinated as shown by anti-TNIP1 immunoblots. Ubiquitinated TNIP1 was stabilized by the addition of concanamycin A (ConA) indicating its lysosomal degradation in treated and nontreated cells. Actin was used as loading control.

B, C Mutations of identified TNIP1 ubiquitination sites do not lead to reduced lysosomal degradation as indicated by stabilized protein amounts by ConA treatment. This is the case for fed control conditions (DMEM) as well as under active autophagy (Rapa and HBSS treatment). (C) shows quantification of blots exemplified in (B) (n=3, error bars indicate std. dev.).

D Mutated TNIP1$^{K371/389R}$ is still getting ubiquitinated as indicated by anti-TNIP1 IP followed by anti-ubiquitin western blot. The addition of ConA leads in all cases to a stabilization of polyubiquitinated protein variants.

**Supplemental Figure S2**

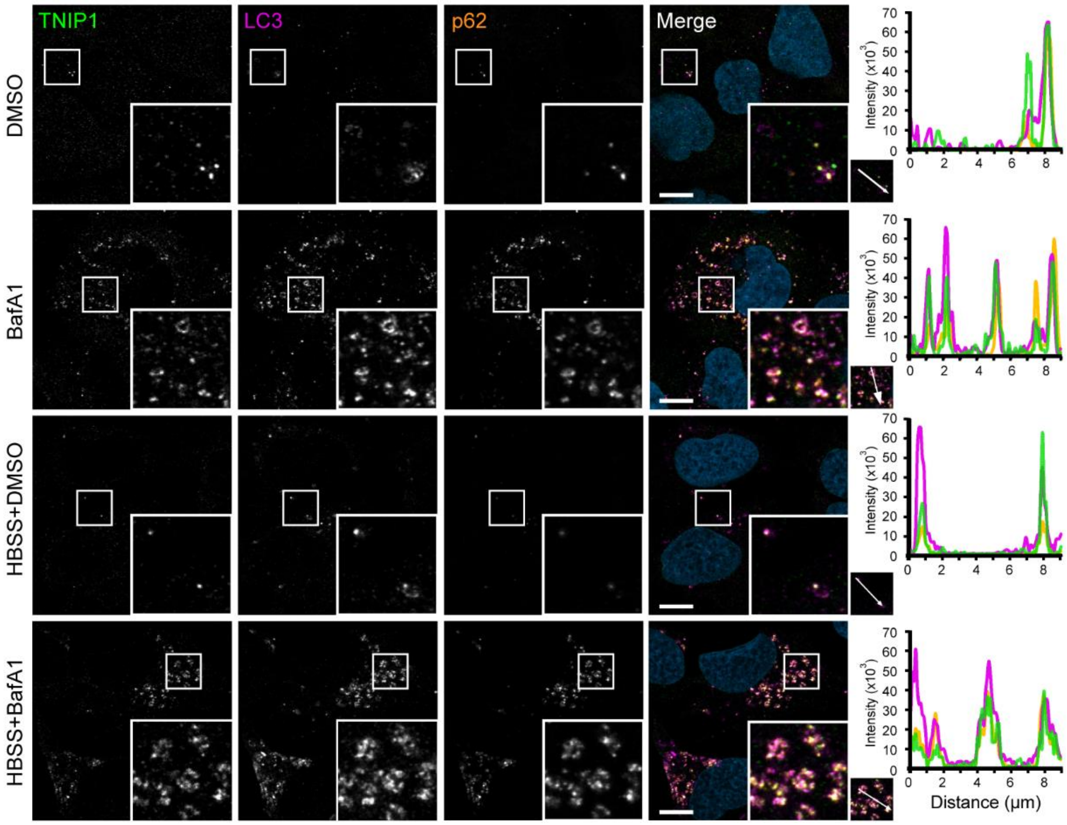


**Supplemental Figure S2. TNIP1 localizes to autophagosomes.**

U2OS cells kept in either fed or starved (HBSS) conditions and treated with either vehicle (DMSO) or BafA1 for 8 hours. Cells were fixed and stained with antibodies against endogenous TNIP1 (green), p62 (orange) and LC3 (purple) and imaged using the Zeiss LSM800 confocal microscope. Line-profile co-localization plots were made using the line-profile quantification tool in the Zen blue imaging software (Zeiss). Vertical axis represents measurements of fluorescent intensity and the horizontal axis the drawn distances. Scale bar = 10 µm.

**Supplemental Figure S3**

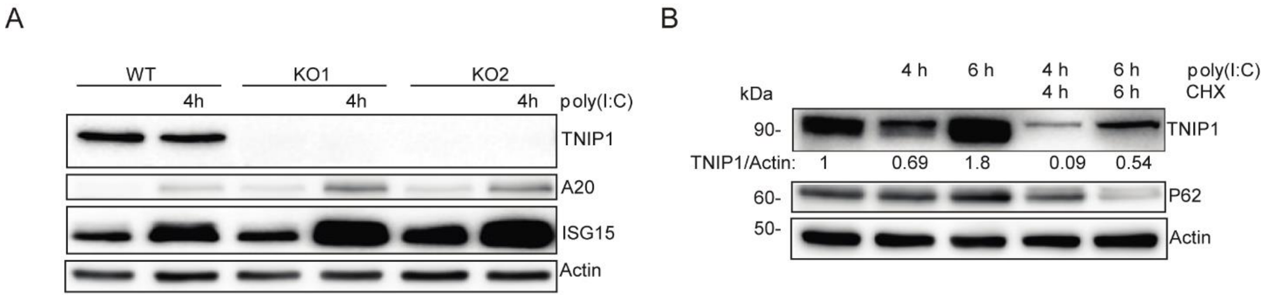


**Supplemental Figure S3. Regulation of TNIP1 protein abundance.**

A Reduction of TNIP1 correlates with an increase of ISG15 and TNFAIP3/A20 in HeLa cells. The increase of the TNIP1 interaction partner TNFAIP3 under poly(I:C) treatment indicates the existence of distinct TNIP1 pools, i.e. free and bound to TNFAIP3.

B Blockage of protein translation by cycloheximide (CHX) treatment reduced the time- and poly(I:C)-dependent increase of TNIP1 after 6 h of treatment indicating a regulation on translational level.
